# A brain-wide form of presynaptic active zone plasticity orchestrates resilience to brain aging in *Drosophila*

**DOI:** 10.1101/2022.06.29.498204

**Authors:** Sheng Huang, Chengji Piao, Christine B. Beuschel, Stephan J. Sigrist

## Abstract

The brain as a central regulator of stress integration determines what is threatening, stores memories and regulates physiological adaptations across the aging trajectory. While sleep homeostasis is linked to brain resilience, how age-associated changes intersect to adapt brain resilience remains enigmatic. We here provide evidence that a brain-wide form of presynaptic active zone plasticity (“PreScale”) promotes resilience by coupling sleep, longevity and memory during aging. PreScale increased until mid-age and contributed to the age-adaption of sleep patterns, in effect promoting longevity but not memory of aging flies. Mechanistically, imaging and electrophysiology suggest that genetically-encoded PreScale reprograms neuronal activity, membrane firing patterns and excitability of the sleep-promoting dorsal fan-shaped body neurons, qualitatively similar to aging. Flies metabolically reprogrammed by spermidine towards extended longevity and preserved memory skipped PreScale and subsequently age-associated sleep pattern changes. Acute deep sleep induction in mid-age flies reset PreScale back to juvenile levels and restored memory. Taken together, early along aging trajectory, PreScale seems to steer trade-offs between longevity and memory, illustrating how life strategy manifests on circuit and synaptic plasticity levels.

**GRAPHIC ABSTRACT:** 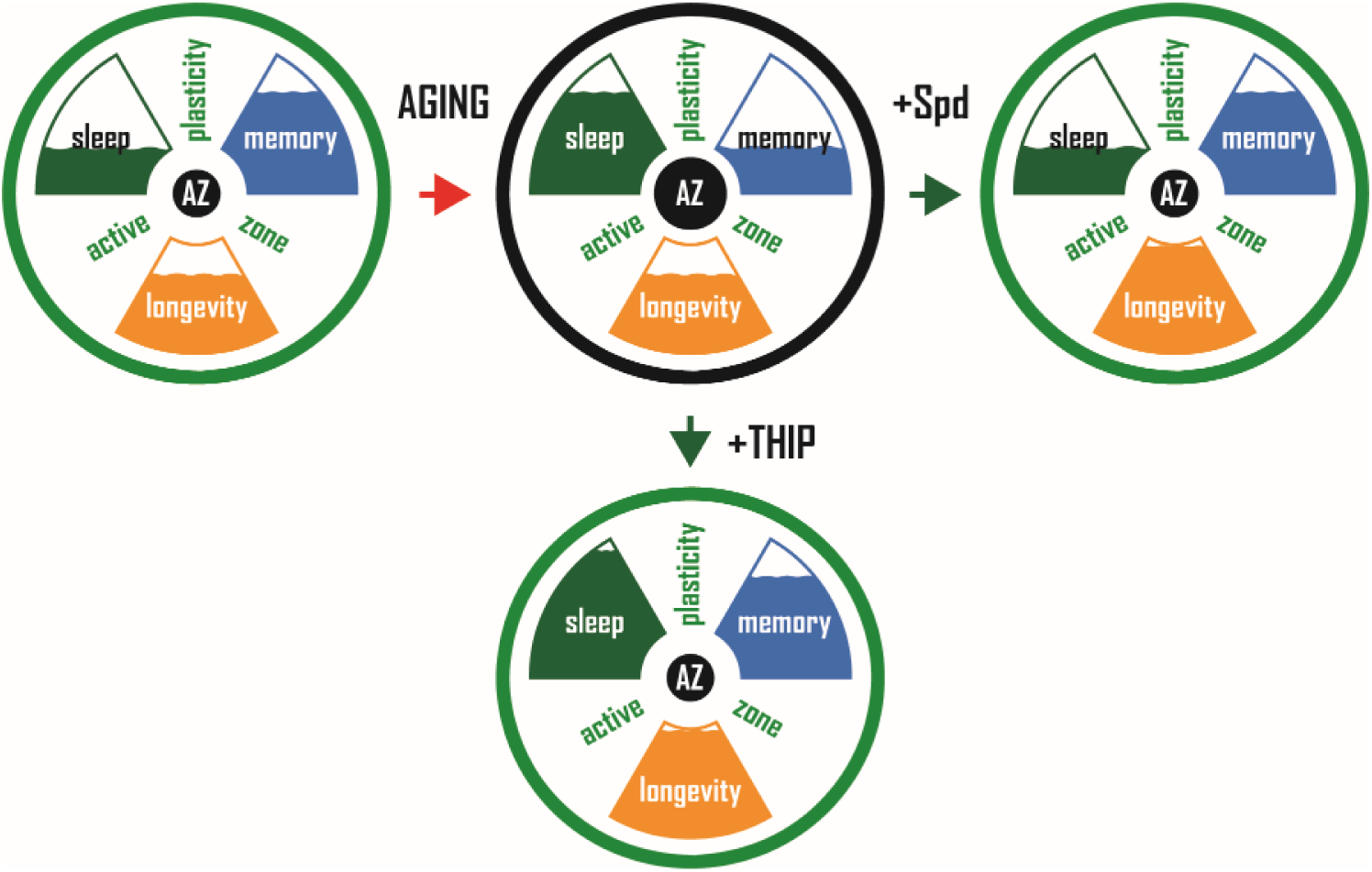

Presynaptic plasticity at the active zone (AZ) during early aging triggers sleep pattern changes, and subsequently steers trade-offs between memory formation and longevity. Interventions like spermidine (Spd) and Gaboxadol (THIP) supplementation suppress PreScale and allow for new memory formation and lifespan extension.

## INTRODUCTION

The aging of the human societies over the globe, with lifespan extension currently being unmatched by increases in healthspan, urges to better understand the molecular-cellular mechanistic underpinnings underlying the various age-associated alterations (Lopez-Otin et al., 2013; Partridge et al., 2018). The current increase of human lifespan expectancy demands for (i) a better understanding of aging mechanisms (Yankner et al., 2008; Bishop et al., 2010; Wang et al., 2011; Morrison and Baxter, 2012; Lopez-Otin et al., 2013; Mander et al., 2013; Gupta et al., 2016; Mander et al., 2017; Partridge et al., 2018) and (ii) safe paradigms to allow for healthspan extension (Eisenberg et al., 2009; Wang et al., 2011; Gupta et al., 2013; Gill et al., 2015; Kaeberlein et al., 2015; Madeo et al., 2018; Tain et al., 2020; Ulgherait et al., 2021). While the resilience against the effects of brain aging is an individual trait (McEwen, 2016; Aron et al., 2022), its representation on cellular and molecular levels remains insufficiently understood. Similarly, the resilience-relevance of age-associated molecular and behavioral changes observed in humans and in animal models remains largely unclear (Bishop et al., 2010; Morrison and Baxter, 2012; Lopez-Otin et al., 2013; Donlea et al., 2014; Mander et al., 2017; Partridge et al., 2018). Specifically, discriminating protective from detrimental changes often remains speculative and unsettled, also impeding the discovery and development of safe healthspan extension paradigms.

Aging provokes changes in both sleep pattern and memory, which are suspected to functionally interact (Donlea et al., 2011; Mander et al., 2013; Mander et al., 2017; Dissel, 2020). The fruitfly *Drosophila* has been widely used in discovering molecular, cellular and circuit mechanisms of memory formation (Quinn and Dudai, 1976; Margulies et al., 2005; Keene and Waddell, 2007), as well as in understanding the mechanisms and functions of sleep (Hendricks et al., 2000; Shaw et al., 2000; Ly et al., 2018). Moreover, *Drosophila* has also been developed into a suitable animal model for studying aging and age-associated sleep pattern alterations (Koh et al., 2006; Vienne et al., 2016; Curran et al., 2019; Ly and Naidoo, 2019; Wiggin et al., 2020) and memory decline (Yamazaki et al., 2007; Gupta et al., 2013). With a relative short lifespan, but not too short, *Drosophila* provides an opportunity to explore and causally connect age-associated molecular and behavioral changes which one day might pave the way for diagnosis and therapy.

We previously described a form of brain-wide presynaptic plasticity (PreScale) during early aging or in reacting to sleep loss, which is characterized by upregulations of synaptic proteins constituting vesicle release sites and accompanied with a decline of efficiency in forming new memories (Gupta et al., 2016; Huang et al., 2020). Notably, rejuvenation paradigm using supplementation of the polyamine spermidine (Spd) attenuates not only the occurrence of age-associated memory decline but also the age-associated emergence of PreScale-type plasticity, while it extends lifespan (Gupta et al., 2016; Tain et al., 2020). However, how exactly PreScale-type plasticity intersects with the several aspects of brain aging, whether in a protective or detrimental manner, remained largely unclear.

We here provide evidence that PreScale operates at the intersection of principal organismal fitness and the formation of new memories during early aging. PreScale peaked at mid-age, changed sleep patterns and promoted lifespan, but at the same time restricted the extent of forming new memories. We speculate that PreScale might be a reversible resilience-conferring plasticity module which reconfigures brain activity patterns to optimize the trade-offs between longevity, memory and sleep during a still plastic phase of early brain aging in *Drosophila*.

## RESULTS

### Global presynaptic active zone plasticity (PreScale) peaks in the mid-aged *Drosophila* brain

In the nervous system, the operation of brain circuits relies on the proper fusion of neurotransmitter-filled synaptic vesicles (SVs) at the so-called presynaptic active zone (Sudhof, 2012; Böhme et al., 2016). Our previous study described a new form of global presynaptic active zone plasticity upon early *Drosophila* aging (Gupta et al., 2016), which here we refer to as PreScale. It is characterized by a brain-wide increase in the ELKS-family core active zone scaffold protein Bruchpilot (BRP) and BRP-associated release factors, particularly (m)Unc13-family protein Unc13A which is crucial for SV release at *Drosophila* synapses (Kittel et al., 2006; Wagh et al., 2006; Böhme et al., 2016; Gupta et al., 2016; Piao and Sigrist, 2022). Notably, BRP seems to operate as a master scaffold of the *Drosophila* active zone which recruits a spectrum of synaptic proteins to tune presynaptic active zone composition and consequently drives synaptic plasticity (Fouquet et al., 2009; Huang et al., 2020; Huang and Sigrist, 2020). Intriguingly, such a global BRP increase associated with a corresponding increase in Unc13A was also observed upon sleep loss (Gilestro et al., 2009; Huang et al., 2020; Huang and Sigrist, 2020). In this context, the extent of sleep loss typically correlates with increases of BRP levels (Gilestro et al., 2009), while genetically enforced sleep reduces BRP levels below baseline levels (Huang et al., 2020).

We here ask for the role of PreScale-type plasticity across the aging trajectory. So far, we had found that BRP/Unc13A levels were upregulated at 30 days (30d) compared to 5 days (5d) of age (Gupta et al., 2016). As the first step, we quantified the levels of BRP and associated synaptic machinery across the *Drosophila* life history. We wondered what limits these presynaptic plastic changes might have and whether BRP would increase gradually across the whole fly lifespan.

Depending on the rearing condition, *Drosophila* can live up to about 3 months which provides an opportunity to explicitly dissect age-associated molecular and behavioral changes (Figure 1A), with the chance to identify critical time windows of turning and switching events. To address our question, we collected brain samples of wildtype (*wt*) at the following ages: 1-day-old (1d), 10d, 20d, 30d, 40d, 50d, 60d and 70d, for quantitative Western blot analysis monitoring a spectrum of pre- and postsynaptic proteins: BRP (Kittel et al., 2006; Wagh et al., 2006), (m)Unc13 family protein Unc13A (Böhme et al., 2016), synaptic vesicle protein Synapsin (Syn) (Klagges et al., 1996), SNARE complex core component Syntaxin (Syx) (Schulze et al., 1995), the homolog of mammalian scaffold protein PSD95 Discs large (Dlg1) (Woods and Bryant, 1991), as well as the crucial autophagy protein Atg8a which is regulated by the aging process (Simonsen et al., 2008). Indeed, BRP, Unc13A, Syn and Dlg1 first increased almost linearly to arrive at a plateau at 30d to 40d (“early aging”) (Figures 1B-1F), while the ratios between activated, lipidated Atg8a (Atg8a-II) and inactive, un-lipidated Atg8a (Atg8a-I) showed a clear trend of gradual reduction during early aging (Figures 1B and 1H), consistent with previous reports (Simonsen et al., 2008; Gupta et al., 2013; Cho et al., 2021). Thereafter, however, the levels of these proteins dropped gradually with further advanced aging (Figures 1A-1F). Syx levels also showed a principally similar behavior (Figures 1B and 1G). While the exact time course might be somewhat variable, an independent experiment with samples covering the phase of early aging using another BRP antibody (Nc82) principally confirmed the age-associated dynamics of synaptic proteins (Figure S1). Consistent with these quantitative Western blot results (Figures 1A-1H), immunostaining identified a relative stronger increase in BRP levels at early aging stages than at later advanced aging stages (Figures 1I and 1J). Non-linear fitting via quadratic regression also suggests maximum BRP, Unc13A, Dlg1 and Syn levels at an age between 30d and 40d in Western blots (Figure 1K). Notably, the dropping of synaptic protein levels at advanced ages, most prominently BRP and Unc13A levels, appeared to be closely associated with declining survival rates (Figures 1A and 1K). These results thus suggest a switch of PreScale at middle age and uncover biphasic synaptic changes across the *Drosophila* lifespan.

**Figure 1.**
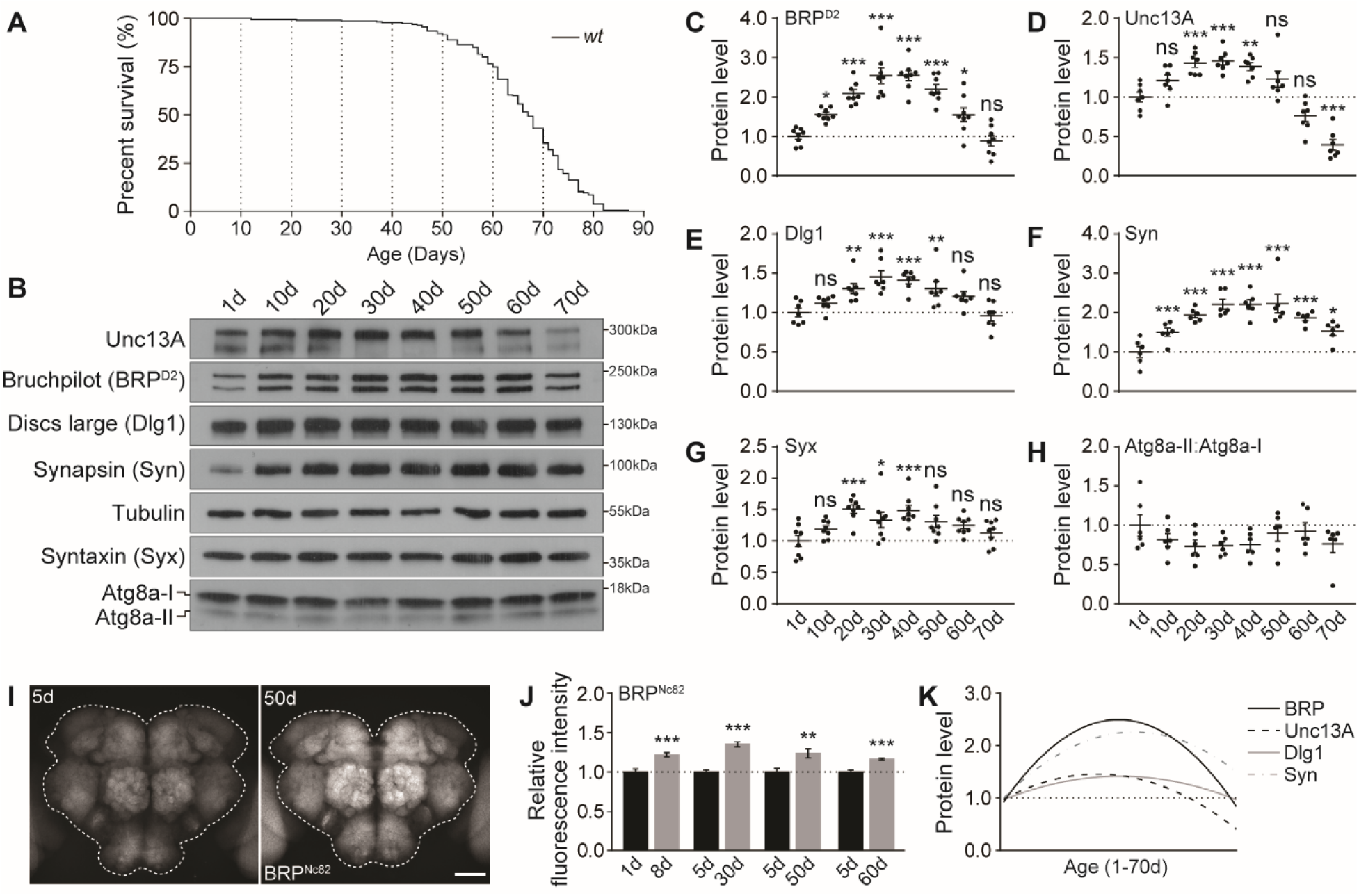
Synaptic plasticity across the fly lifespan. (**A**) A typical survival curve of *w^1118^ wt* female flies. n = 235. (**B**-**H**) Representative Western blots (**B**) and relative levels of a spectrum of synaptic proteins, including BRP (**C**), Unc13A (**D**), Dlg1 (**E**), Syn (**F**), and Syx (**G**), and the relative ratios of active lipidated and un-lipidated autophagy related protein Atg8a (Atg8a-II:Atg8-I, **H**) in *wt* flies across the lifespan. n = 6-8. One-way ANOVA with Dunnett’s post hoc tests is shown. (**I** and **J**) Confocal images (**I**) and whole-mount brain staining analysis (**J**) of BRP in *wt* flies at different ages. n = 15 for 8d, n = 9 for 30d, n = 14-15 for 50d and n = 9 for 60d. Mann-Whitney test is shown. *p < 0.05; **p < 0.01; ***p < 0.001; ns, not significant. Scale bar: 50 μm. Error bars: mean ± SEM. (**K**) Non-linear line fitting quadratic regression of the protein levels of BRP (**C**), Unc13A (**D**), Dlg1 (**E**), Syn (**F**) levels in Western blot across the fly lifespan.

### PreScale dampens both spontaneous activity and membrane excitability of sleep-controlling dFB neurons

What is the circuit and brain activity relevance of the BRP-driven synaptic plasticity we call PreScale? As the master scaffold of the active zone, BRP controls the synaptic vesicle (SV) release properties, e.g., release probability and/or release site number (Kittel et al., 2006; Böhme et al., 2016; Huang et al., 2020). Concerning changes of synaptic transmission in response to changing BRP levels, recent analysis at the physiologically highly accessible larval neuromuscular junction comparing 4xBRP to 2xBRP control animals showed largely normal synaptic currents when evoked by single discrete action potentials (Huang et al., 2020). However, short-term plasticity as well as the evoked transmission upon 10 Hz stimulation were markedly facilitated in 4xBRP animals (Figure S4M in (Huang et al., 2020)). Thus, PreScale might promote the transmission of specific activity patterns in certain frequency domains, for example the oscillatory activity pattern changes in the R5 network of sleep-deprived animals (Raccuglia et al., 2019).

Indeed, we previously showed that 4xBRP could mimic mid-aged animals in age-associated memory decline (Gupta et al., 2016). Given that an increase of BRP copy number encodes sleep need (Huang et al., 2020), we recently showed an increased activity level in the sleep-promoting R5 neurons of the ellipsoid body by using the Ca^2+^-dependent activity reporter CaLexA (Figures S2A and S2B) (Masuyama et al., 2012; Liu et al., 2016; Huang et al., 2020). To further analyze in a brain-wide manner, CaLexA was expressed by the pan-neuronal driver *elav-Gal4*. The expression pattern of *elav-Gal4* is known to be strong in antennal lobe (AL) and mushroom body (MB), and indeed *elav-Gal4*- driven CaLexA showed strong signals in AL and MB (Figure 2A). Interestingly, in 4xBRP animals CaLexA expression within the ALs was significantly reduced while staying largely unaffected in MBs (Figures 2A and 2B). As the AL operates as the first synaptic relay station of the olfactory system and a central gate of olfactory sensory information processing (Davis, 2011), a generic drop in its activity levels might potentially support sleep. In another experiment, we expressed CaLexA in the well-studied sleep-promoting dorsal fan-shaped body (dFB) using *R23E10-Gal4* drivers (Donlea et al., 2011; Pimentel et al., 2016). dFB neurons express allatostatin-A peptide that inhibits helicon cells which innervate and activate R5 neurons (Donlea et al., 2018). Consistent with the inhibitory connection between dFB and R5, we observed a strong activity decrease in dFB in 3xBRP and 4xBRP animals (Figures 2C and 2D), while the activity of R5 was significantly increased (Figures S2A and S2B) (Huang et al., 2020).

**Figure 2.**
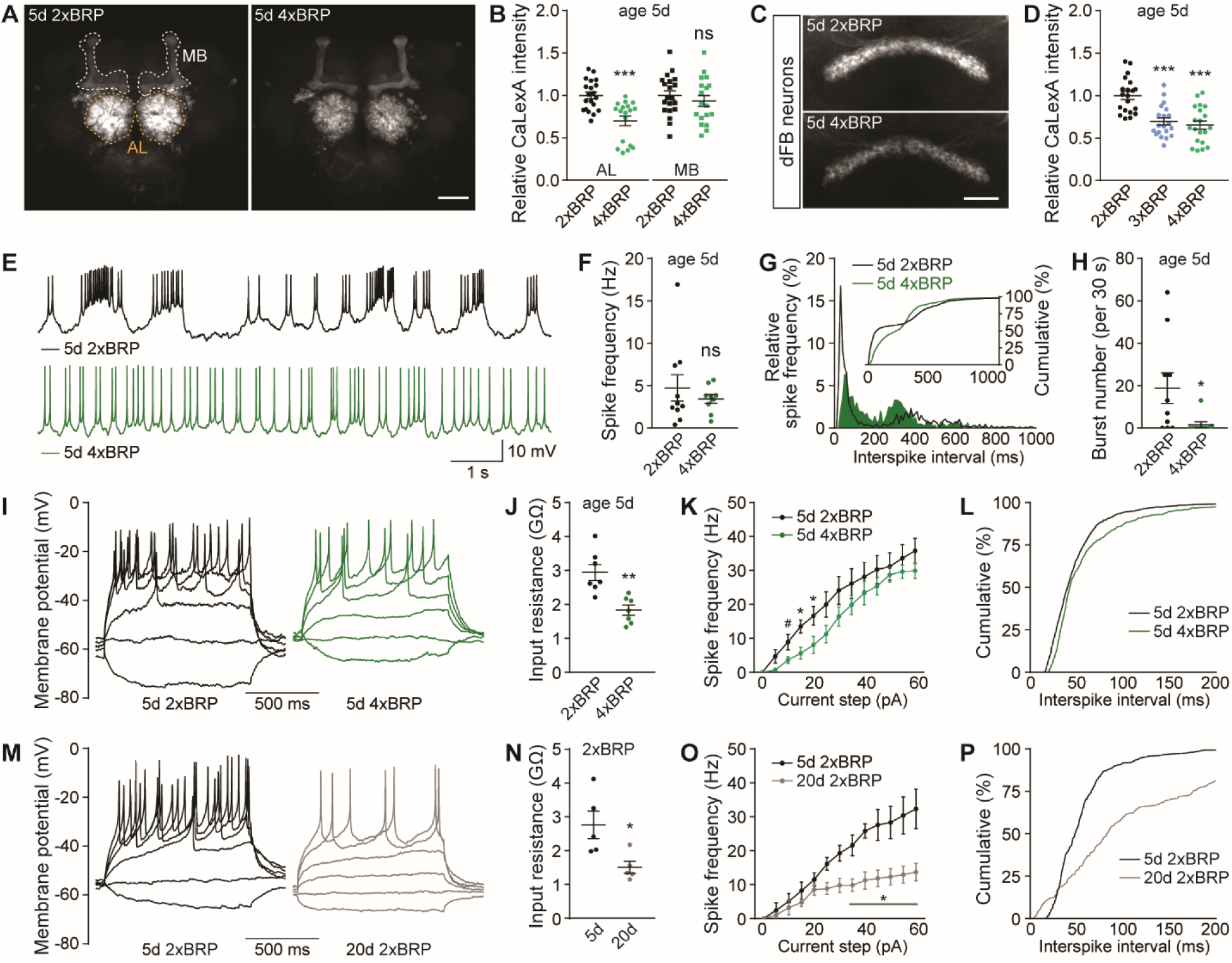
PreScale plasticity provokes activity reprogramming in the dorsal fan-shaped body (dFB) neurons. (**A** and **B**) Confocal images (**A**) and whole-mount brain staining analysis (**B**) of CaLexA signal intensity with CaLexA expressed pan-neuronally by *elav-Gal4* in 2xBRP compared to 4xBRP female flies. n = 18-20 for all groups. MB, mushroom body, AL, antennal lobe. Scale bar: 50 μm. (**C** and **D**) Confocal images (**C**) and whole-mount brain staining analysis (**D**) of CaLexA signal intensity with CaLexA expressed in dorsal fan-shaped body (dFB) by *R23E10-Gal4* in 2xBRP compared to 3xBRP and 4xBRP female flies. n = 20 for all groups. Scale bar: 20 μm. (**E**-**H**) Spontaneous firing of dFB neurons in 5d 2xBRP and 4xBRP flies, including representative membrane potential traces (**E**), mean firing rate (**F**), interspike interval distributions (**G**, Kolmogorov-Smirnov test, p < 0.0001), and bursting number for a 30s window (**H**, Mann-Whitney test, p = 0.0212). n = 9-10 per group. (**I**-**L**) Voltage responses to 1-s current steps (from -10 to 60pA, 5pA) in dFB of 5d 2xBRP and 4xBRP flies, including representative traces (**I**), input resistances (**J**), firing rates (**K**, two-way repeated-measures ANOVA analysis, p = 0.0245 for Current × Genotype interaction), and distribution of interspike interval (**L**, Kolmogorov-Smirnov test, p < 0.0001). (**M**-**P**) Voltage responses to 1-s current steps (from -10 to 60pA, 5pA) in dFB of 5d and 20d 2xBRP flies, including representative traces (**M**), input resistances (**N**), firing rates (**O**, two-way repeated-measures ANOVA analysis, p < 0.0001 for Current × Age interaction), and distribution of interspike interval (**P**, Kolmogorov-Smirnov test, p < 0.0001). Mann Whitney test is shown for comparison between two groups. One-way ANOVA with Tukey’s post hoc tests is shown for comparisons of three groups. #p = 0.05; *p < 0.05; ** p < 0.01; *** p < 0.001; ns, not significant. Error bars: mean ± SEM.

As suggested by the CaLexA results in dFB at age 5d genetically triggered by PreScale (Figures 2C and 2D) and the well-characterized role of the dFB neurons in promoting sleep (Donlea et al., 2011; Pimentel et al., 2016), we speculated that the plastic remodeling of neuronal activity and firing patterns in dFB neurons might be changed upon early aging. Moreover, if PreScale was really responsible to age-adapt dFB physiology, it might be possible to impose similar dFB changes in young animals through genetically installing PreScale. Thus, we performed current clamp of the dFB neurons (Figures S2C and S2D), measured their spontaneous spiking patterns and injected current steps to elicit membrane voltage responses. Notably, though mean spontaneous spiking activity frequency remained unaffected in 4xBRP animals, the regularity of spontaneous spiking was enhanced and bursting spikes were rarely seen in 4xBRP dFB neurons (Figures 2E-2H). Consistent with a recent study showing that the regularity of spontaneous firing of the sleep-regulating clock DN1p neurons determines sleep quality (Tabuchi et al., 2018), the pattern changes of spontaneous firing of dFB neurons might contribute to sleep regulation in 4xBRP animals.

To determine the membrane excitability properties of dFB neurons comparing 4xBRP with 2xBRP, we injected current steps to depolarize the membrane and elicit action potentials. The dFB neurons in 4xBRP flies tended to be electrically silent, with lower input resistances, fewer action potentials elicited and increased interspike intervals (Figures 2I-2L), in agreement with the CaLexA results (Figures 2C and 2D). To directly compare the effects of PreScale to potential effects of aging here, we intended to measure the electric properties of dFB at an age of 30d. Unfortunately, however, rates of successful patching were extremely low in these aged animals. However, at age 20d, we could successfully measure a few dFB neurons (Figures 2M-2P). The dFB neuron were indeed more electric silent, which showed reduced input resistances and increased interspike intervals, and current step elicited action potential firing almost failed to raise in 20d flies. While qualitatively similar, effect sizes were clearly more pronounced in the comparison of 20d to 5d than comparing 2xBRP to 4xBRP at age 5d. These data suggest that the genetically installed PreScale can operate as surrogate for important aspects of early aging and interventions of PreScale-type plasticity might be potential paradigms for healthy aging.

Taken together, our imaging and electrophysiology results point towards a reprograming of neuronal activity across the fly brain under PreScale-type plasticity, which likely reflects a shared synaptic origin of sleep homeostasis and early brain aging. The functional reprograming of the sleep-promoting dFB and R5 network under PreScale suggests that PreScale might reprogram early aging-associated behavioral changes. We went on testing this hypothesis.

### PreScale promotes survival over memory formation during early aging

In our previous study, as also mentioned earlier, we could linearly increase BRP levels by titrating the number of *brp* genomic copies, which promotes sleep in a dosage-dependent manner (Figures S3A-S3F) (Huang et al., 2020), and mimics essential aspects of brain aging (Gupta et al., 2016). Direct quantitative comparisons of BRP levels between early aging and genetic *brp* titration at age 5d by line-fitting indicate that *per se* both scenarios provoked BRP upregulations to similar extents (Figure 3A). If establishing certain BRP and the associated changes in Un13A levels were of specific functions, especially throughout early aging stages, genetically titrating BRP levels might be able to uncover mechanisms of age-associated behavioral changes.

**Figure 3.**
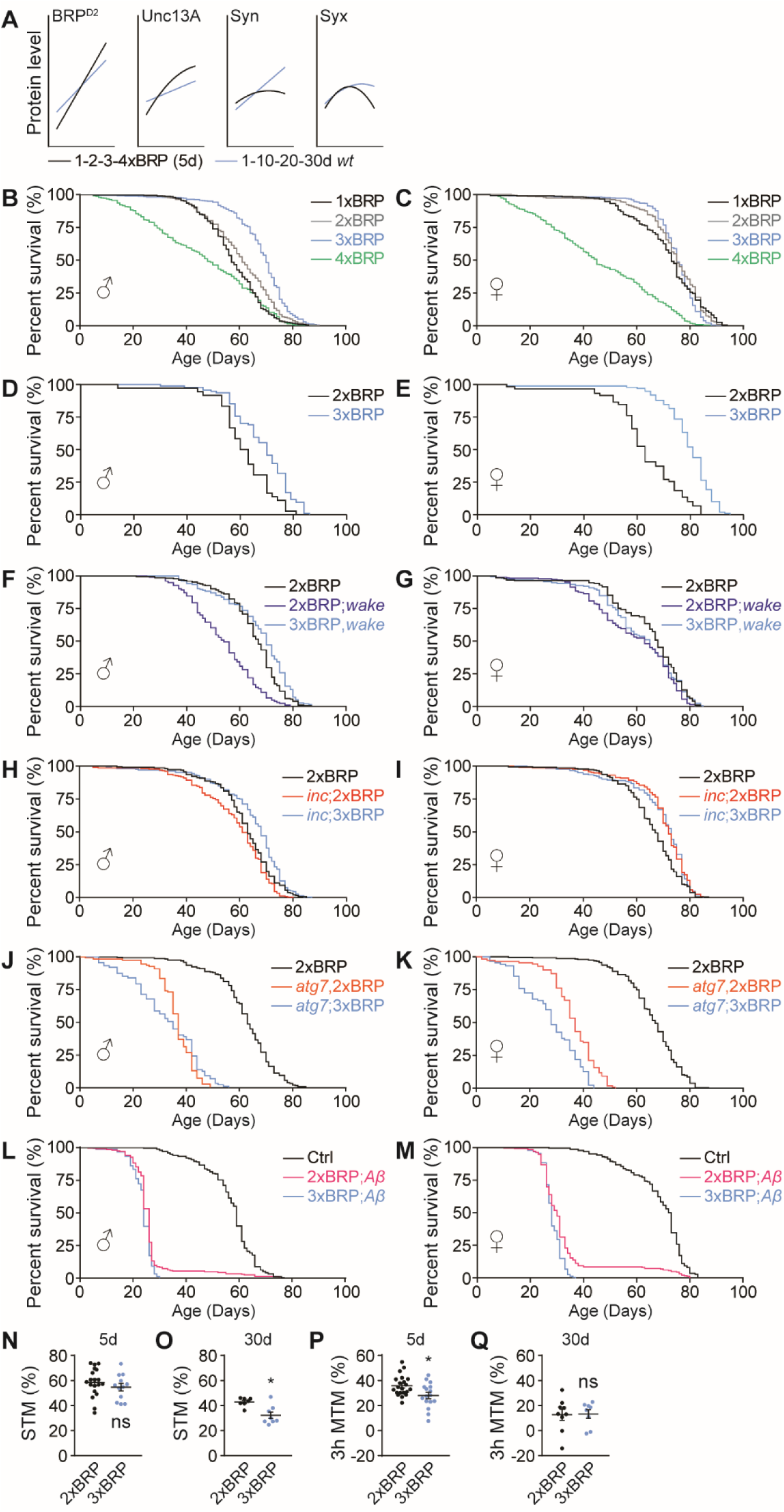
PreScale promotes survival over memory formation during early aging. (**A**) Direct line-fitting comparisons between PreScale during early-stage aging from 1d to 30d (Figure 1) and genetic *brp* titration from 1xBRP to 4xBRP at age 5d (Huang et al., 2020). (**B** and **C**) Lifespan analysis of flies with BRP titration from 1xBRP to 4xBRP. For male flies (**B**), n = 394 for 1xBRP (p < 0.001), n = 396 for 3xBRP (p < 0.001) and n = 395 for 4xBRP (p < 0.001), compared to 2xBRP *wt* control flies (n = 393). For female flies (**C**), n = 400 for 1xBRP (p = 0.017), n = 399 for 3xBRP (ns) and n = 397 for 4xBRP (p < 0.001), compared to 2xBRP *wt* control flies (n = 399). (**D** and **E**) An independent experiment of the lifespan of 2xBRP and 3xBRP flies. For male flies (**D**), n = 94 for 3xBRP compared to 2xBRP (n = 36, p < 0.001). For female flies (**E**), n = 98 for 3xBRP compared to 2xBRP *wt* control flies (n = 59, p < 0.001). (**F** and **G**) Lifespan analysis of flies with 3xBRP in *wake* mutant background. For male flies (**F**), n = 234 for 2xBRP;*wake* compared to 2xBRP *wt* control (n = 233, p < 0.001) or compared to 3xBRP,*wake* (n = 155, p < 0.001). For female flies (**G**), n = 235 for 2xBRP;*wake* compared to 2xBRP *wt* control (n = 230, p < 0.001) or compared to 3xBRP,*wake* (n = 231, ns). (**H** and **I**) Lifespan analysis of flies with 3xBRP in *inc* mutant background. For male flies (**H**), n = 236 for *inc*;2xBRP compared to 2xBRP *wt* control (n = 232, p = 0.009) or compared to *inc*;3xBRP (n = 212, p < 0.001). For female flies (**I**), n = 235 for *inc*;2xBRP compared to 2xBRP *wt* control (n = 235, p < 0.001, *wt* is also shown in Figure 1A) or compared to *inc*;3xBRP (n = 182, ns). (**J** and **K**) Lifespan analysis of flies with 3xBRP in *atg7* mutant background. For male flies (**J**), n = 108 for *atg7*,2xBRP compared to 2xBRP *wt* control (n = 232, p < 0.001) or compared to *atg7*;3xBRP (n = 87, ns). For female flies (**K**), n = 109 for *atg7*,2xBRP compared to 2xBRP *wt* control (n = 235, p < 0.001) or compared to *atg7*;3xBRP (n = 87, p < 0.001). (**L** and **M**) Lifespan analysis of flies with 3xBRP in Alzheimer’s disease model flies (*elav>Aβ*). For male flies (**L**), n = 203 for 2xBRP;*Aβ* compared to *elav>mCD8-GFP* control flies (n = 238, p < 0.001) or compared to 3xBRP;*Aβ* (n = 151, p = 0.006). For female flies (**M**), n = 249 for 2xBRP;*Aβ* compared to *elav>mCD8-GFP* control flies (n = 228, p < 0.001) or compared to 3xBRP;*Aβ* (n = 241, p = 0.002). Gehan-Breslow-Wilcoxon test with Bonferroni correction for multiple comparisons is shown for all longevity experiments. (**N** and **O**) Short-term memory (STM) for 5d (**N**) and 30d (**O**) 2xBRP and 3xBRP flies. n = 19 for 5d 2xBRP, n = 12 for 5d 3xBRP. n = 7-8 for 30d for both groups at 30d. (**P** and **Q**) Middle-term memory (MTM) tested 3 hours after training for 5d (**P**) and 30d (**Q**) 2xBRP and 3xBRP flies. n = 21 for 5d 2xBRP, n = 16 for 5d 3xBRP. n = 8-9 for 30d for both groups at 30d. Mann-Whitney test is shown. *p < 0.05; ns, not significant. Error bars: mean ± SEM.

To this end, we first monitored the longevity of flies with titrated BRP levels and observed an extension of lifespan when increasing *brp* copies stepwise from 1 to 3. However, the longevity of 4xBRP suffered severely, with lifespan clearly shorter than in 2xBRP *wt* animals (Figures 3B-3E and S3G-S3Q). Thus, it is tempting to speculate that a specific BRP level increase might be established to protect the survival of animals from stress conditions, for example in response to sleep deprivation (Gilestro et al., 2009; Huang et al., 2020; Huang and Sigrist, 2020; Weiss and Donlea, 2021) and during early brain aging (Figure 1).

If moderately elevated PreScale was indeed protective for survival, it might be particularly protective for flies challenged by chronic sleep loss and/or flies suffered from reduced lifespan. Thus, we established a 3xBRP situation in two sleep mutants, *wide awake* (*wake*) and *insomniac* (*inc*) mutants with reduced sleep and shortened lifespan (Stavropoulos and Young, 2011; Pfeiffenberger and Allada, 2012; Liu et al., 2014), which have been recently shown to trigger PreScale likely in response to their sleep loss (Huang et al., 2020). Indeed, 3xBRP rescued the lifespan of *wake* and *inc* mutants back to *wt* level (Figures 3F-3I). Interestingly, the lifespan of *inc* female flies was longer than *wt* and not further extended in 3xBRP background (Figure 3I). However, the survival of *atg7* autophagy mutant as well as of Alzheimer’s disease model flies was not protected (Figures 3J-3M), supporting a specific, context-dependent role of PreScale in regulating lifespan.

As robustly observed in the aging process of several model organisms and humans, complex behaviors like memory formation and consolidation typically dwindle in their efficiency with advancing age (Buckner, 2004; Saitoe et al., 2005; Wang et al., 2011). Healthy aging, as an emerging concept (Kaeberlein et al., 2015; Madeo et al., 2018), demands for not only longer life expectance but also specifically for a maintenance of life quality, importantly proper locomotor ability, as well as efficient sleep and memories at older ages. Thus, we next asked whether 3xBRP might be able to delay the normal age-associated memory decline, which has been broadly reported for *Drosophila* (Saitoe et al., 2005; Yamazaki et al., 2007; Gupta et al., 2016). Somewhat surprisingly, however, 3xBRP animals, while showing normal short-term memory (STM) at age 5d, displayed a significant reduction in STM at age 30d (Figures 3N and 3O). Middle-term aversive olfactory memory scores tested 3 hours after training (3h MTM) were already significantly reduced in 3xBRP animals at age 5d (Figures 3P and 3Q).

Thus, moderately elevated PreScale, while promoting lifespan in a context-dependent manner, at the same time did not age-protect memory formation. Given the dynamic nature of PreScale across the fly lifespan (Figures 1 and S1) and the lifespan extension of 3xBRP flies (Figures 3B-3E), PreScale might execute trade-offs to favor survival over complex and costly behaviors like forming new memories (Placais and Preat, 2013; Placais et al., 2017).

### PreScale mediates early aging-associated alterations in sleep behavior

An aging-associated decline of daily locomotor activities is typical for animal models and humans (Carey et al., 2006; Marck et al., 2017), which is often accompanied by sleep pattern changes (Koh et al., 2006; Metaxakis et al., 2014; Wiggin et al., 2020). Furthermore, PreScale seems to reprogram the activity of the interconnected R5/dFB network (Figures 2C-2P and S2). If the age-associated BRP increase was causally connected to locomotor and sleep pattern changes, genetic attenuation of PreScale might result in a delayed onset of locomotor decline and sleep pattern changes during early aging.

We recently showed that removing an endogenous *brp* gene copy out of two by introducing a null mutation in the *brp* locus (referred to as 1xBRP) significantly reduces BRP levels and subsequently provokes a downregulation of other crucial synaptic proteins, importantly Unc13A as well as Syntaxin (Huang et al., 2020). Interestingly, we also observed that while 1xBRP animals displayed only slightly reduced total sleep under baseline condition, they showed facilitated wakefulness under mechanical disturbance at age 5d (Huang et al., 2020). Here, we examined the daily locomotor activity and sleep patterns across early aging stages from age 5d over 20d and 30d to an age of 40d in 1xBRP compared to 2xBRP.

While a robust gradual reduction of total daily locomotor activity was evident for both 2xBRP control and 1xBRP flies during early aging, 1xBRP exhibited higher daily locomotor activity compared to 2xBRP animals across all ages tested (Figures 4A and 4B). This difference was most prominent at age 20d during daytime, especially at the light/dark transition at ZT12 (zeitgeber time) across the tested ages (Figures 4A and 4B). Notably, 1xBRP protracted early aging-associated changes relative to 2xBRP flies in the pattern and amount of daily locomotor activity, evident in 20d 1xBRP resembling 5d 2xBRP, 30d 1xBRP being similar to 20d 2xBRP flies, and so on (Figures 4A and 4B). In short, it seems that during early aging, 1xBRP animals mimicked younger 2xBRP control flies concerning total locomotor activity. Such an effect was even more pronounced in early aging-associated alterations in sleep pattern, especially daytime sleepiness, which was clearly suppressed in 1xBRP animals at all the ages tested (Figures 4C-4G). The suppression of daytime sleepiness was stronger at younger ages (i.e., 20d) evident in a substantial reduction of sleep episode number and duration (Figures 4C-4F). Of note, protraction of the decrease of sleep latency (at ZT0) was also observed for 1xBRP animals (Figure 4G). Nighttime sleep was also significantly suppressed by 1xBRP at different ages compared with age-matched 2xBRP control flies (Figure 4H). Interestingly, nighttime sleep was less consolidated and the difference in sleep latency at ZT12 diminished with aging in 1xBRP animals compared to 2xBRP flies (Figures 4I-4K).

**Figure 4.**
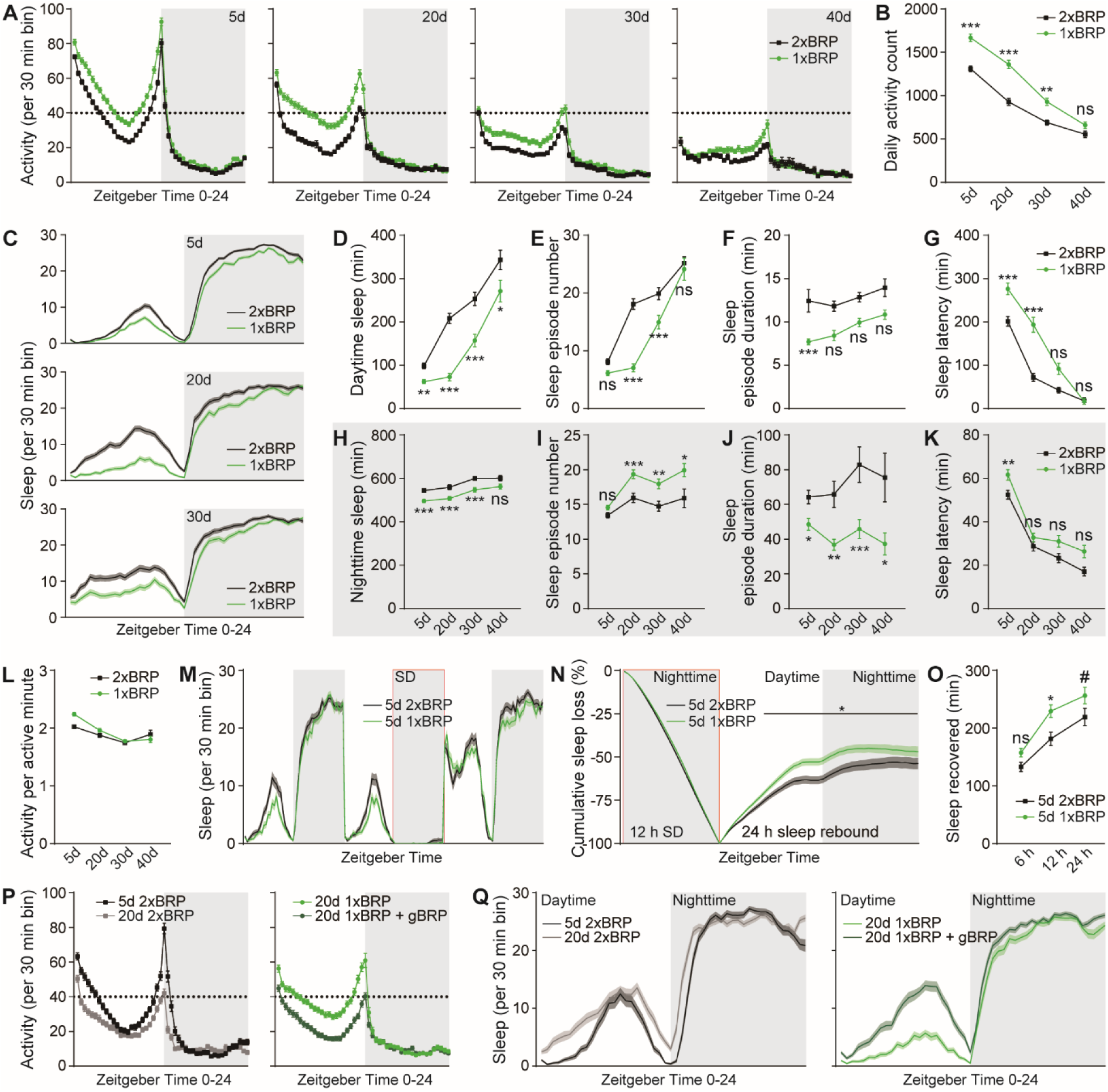
PreScale mediates early aging-associated sleep pattern changes. (**A** and **B**) Locomotor walking activity distribution across the day (**A**) and averaged daily total walking activity (**B**) of 1xBRP compared to 2xBRP at ages 5d, 20d, 30d and 40d. (**C**-**K**) Sleep structure of 1xBRP and 2xBRP flies at ages 5d, 20d, 30d and 40d averaged from measurements over 2-3 days, including sleep profile plotted in 30-min bins (**C**), daytime and nighttime sleep amount (**D** and **H**), number and duration of sleep episodes (**E**, **F**, **I** and **J**), and sleep latencies (**G** and **K**). n = 190-191 for 5d, n = 94-95 for 20d, n = 77-78 for 30d and n = 32 for 40d. Two-way ANOVA with Sidak’s multiple comparisons is shown. (**L**) Activity index (average activity count per active minute) of 1xBRP and 2xBRP animals at ages 5d, 20d, 30d and 40d. Correlation analysis did not identify any significant age-associated difference in activity index of both 1xBRP and 2xBRP animals. (**M**) Sleep profile for 2xBRP *wt* control and 1xBRP flies for 3 consecutive days. SD, sleep deprivation. (**N**) Normalized cumulative sleep loss during 12 h nighttime sleep deprivation and 24 h sleep rebound. n = 116-121 for both groups. Two-way repeated-measures ANOVA with Fisher’s least significant difference (LSD) test detected a significant genotype × time interaction (F_(47, 11233)_ = 3.933; p < 0.0001) during sleep rebound. Asterisks above a line indicate time points at which the sleep percentage recovered differs significantly between female 2xBRP *wt* and 1xBRP flies, respectively. (**O**) Sleep recovered at three different time points after sleep deprivation for female 1xBRP compared to 2xBRP *wt* flies at age 5d. Two-way ANOVA with Sidak’s multiple comparisons is shown. (**P** and **Q**) Locomotor walking activity (**P**) and sleep profile (**Q**) of 1xBRP flies at age 20d rescued by introducing a transgenic *brp* copy (gBRP, *brp* P[acman]). n = 63-64 for all groups. *p < 0.05; **p < 0.01; ***p < 0.001; ns, not significant; #p = 0.07. Error bars: mean ± SEM.

We speculated that if the 1xBRP constellation would attenuate the early aging-associated changes of sleep pattern, it might also show stronger sleep rebound upon sleep deprivation, as aging is typically associated with a decline in sleep rebound in response to sleep deprivation (Vienne et al., 2016). To test this idea, we sleep-deprived both 1xBRP and 2xBRP flies at age 5d during nighttime for measuring sleep rebound afterwards (Figure 4M). In agreement with our speculation, 1xBRP flies indeed exhibited enhanced sleep rebound (Figures 4N and 4O). We here also performed an additional control experiment by putting an artificial genomic *brp* gene copy back to 1xBRP flies which should restore *wt*-type sleep patterns. Indeed, the early aging-associated sleep pattern changes normally seen in 2xBRP *wt* flies were restored (Figures 4P, 4Q and S4A-S4D), further supporting the specific role of BRP-driven PreScale-type plasticity in promoting early aging-associated sleep pattern changes and survival. At the first glance, an apparent paradox emerges in that 1xBRP animals concerning their sleep pattern regulation appear to age slower but are ultimately shorter-lived, when compared to 2xBRP controls (Figures 3B, 3C and 4). Together with the extended lifespan of 3xBRP animals (Figures 3B-3E), this might be explained by the early aging-associated alterations of locomotor activity and sleep behavior being age-adaptive and survival-protective (Figures 4D-4L).

Contrary to 1xBRP (Figure 4), 3xBRP likely operates as a mimicry of the essential aspects of brain aging, which favors survival over cognitive functions (Figures 1, S1, 2C-2P, 3 and S3). Indeed, 3xBRP animals displayed a mild reduction in total locomotor activity (Figures S4E and S4F) and increased and consolidated sleep (Figures S4G-S4O) compared to 2xBRP controls at age 5d. These differences diminished gradually with aging (Figures S4E-S4O). It appears that the genetically installed moderate PreScale in 3xBRP animals likely augments an early aging-associated brain reprogramming, which is essential for later survival during advanced aging, as e.g. also suggested by the consequences of sleep loss early in life (Seugnet et al., 2011; Kayser et al., 2014).

### Spermidine supplementation attenuates early aging-associated sleep pattern changes

Aging is typically associated with behavioral alterations affecting life quality and consequent burden of the aging human societies (Kaeberlein et al., 2015; Madeo et al., 2018). Spermidine (Spd) supplementation has recently emerged as a promising paradigm to promote healthy aging, supported by studies ranging from flies and rodent models to human clinical trials (Madeo et al., 2018). In flies, Spd supplementation was shown to increase lifespan and attenuate early aging-associated memory decline (Gupta et al., 2013; Tain et al., 2020; Schroeder et al., 2021). Spermidine supplementation effects are based on metabolic alterations, age-protecting autophagic clearance and mitochondrial respiration (Gupta et al., 2013; Tain et al., 2020; Liang et al., 2021; Schroeder et al., 2021). Importantly, we previously showed that the age-associated PreScale is efficiently suppressed by Spd supplementation during early aging stages (e.g. at age 30d) while it does not affect BRP levels in young animals (Gupta et al., 2016). These studies, together with our current findings (Figures 1 and S1), suggest that Spd supplementation should be able to delay the onset of age-associated behavioral alterations, for example locomotion and sleep pattern (Figure S5A).

Indeed, the daily locomotor activity of Spd-supplied 30d flies was significantly higher when compared with age-matched control flies, while no difference was observed in 5d young animals (Figures 5A and 5B). Moreover, the early aging-associated sleep pattern changes were suppressed to a certain degree, especially concerning the daytime sleep phenotypes (Figures 5C-5G). Here, daytime sleep increases were substantially suppressed through a significant reduction in the number of sleep episodes with concomitant increases of sleep latency (Figures 5C-5E and 5G). Nighttime sleep changes were also suppressed mainly through an increase of sleep latency towards the values normally found in 5d young flies, while the number of nighttime sleep episodes and mean sleep episode duration stayed comparable to unsupplied siblings (Figures 5C-5G). All these changes were specific for 30d flies, as the sleep of 5d young flies was not affected by Spd supplementation (Figures 5A-5G), consistent with the notion that Spd supplementation specifically affects the aging process of flies (Gupta et al., 2013; Gupta et al., 2016; Liang et al., 2021; Schroeder et al., 2021). Importantly, the typical age-associated reduction of sleep rebound upon sleep deprivation (Figures 5H-5J) (Vienne et al., 2016) was suppressed as well, allowing for a more efficient sleep compensation when reacting to sleep loss in 30d flies (Figures 5K-5M), but not for 5d flies (Figures S5B-S5E).

**Figure 5.**
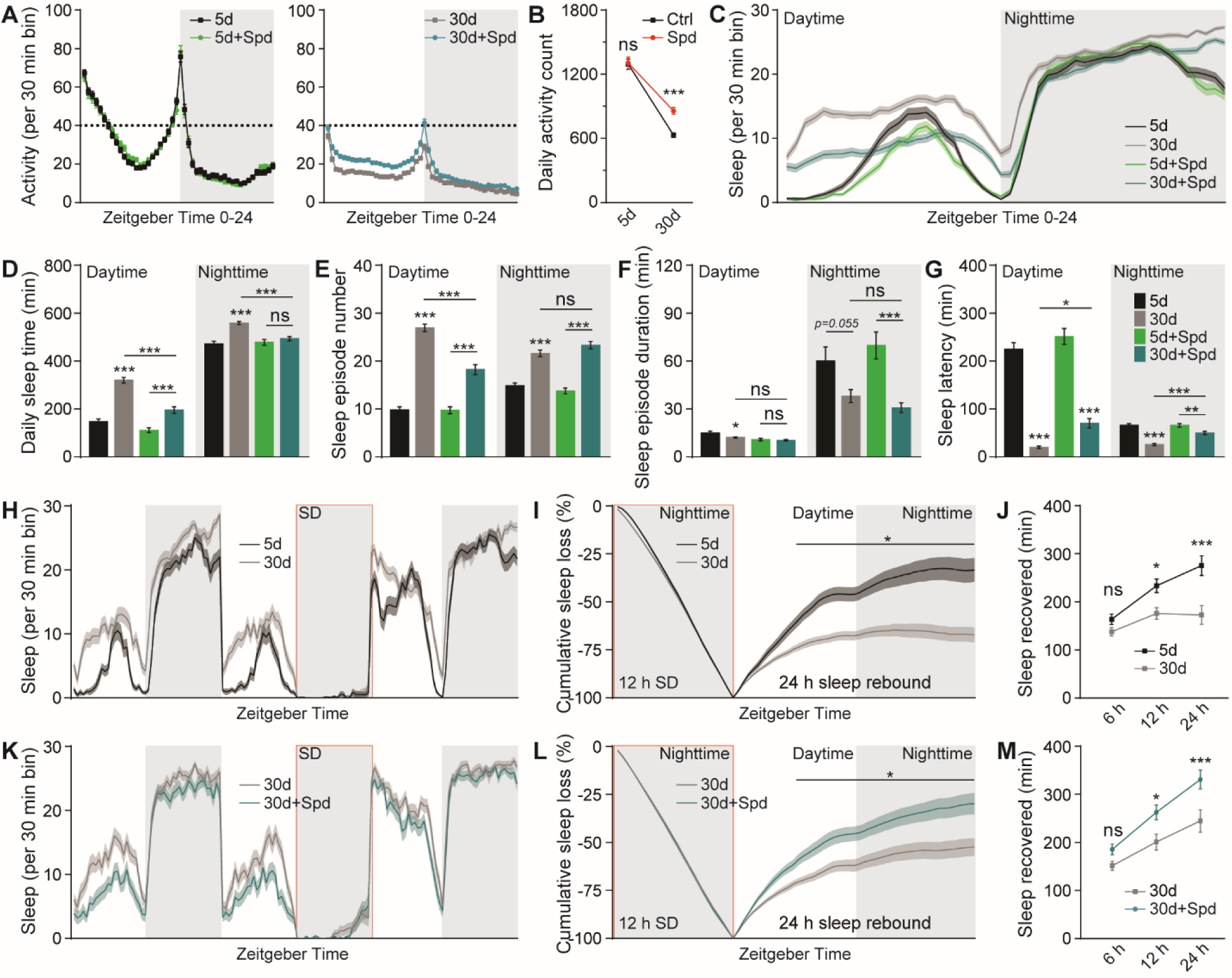
Spermidine supplementation attenuates early aging-associated sleep pattern changes. (**A**) Locomotor walking activity of 5 mM Spd-treated *wt* flies across the day at ages 5d and 30d. (**B**) Averaged daily total locomotor walking activity counts of 5 mM Spd-treated *wt* flies at ages 5d and 30d. Two-way ANOVA Ordinary test is shown. (**C**-**G**) Sleep structure of 5 mM Spd-treated *wt* flies at ages 5d and 30d averaged from measurements over 2-3 days, including sleep profile plotted in 30-min bins (**C**), daytime and nighttime sleep amount (**D**), number and duration of sleep episodes (**E** and **F**), and sleep latencies (**G**). n = 93-96 for all groups. One-way ANOVA with Tukey’s post hoc tests is shown. (**H**) Sleep profile for *wt* flies at age 30d compared to 5d for 3 consecutive days. (**I**) Normalized cumulative sleep loss during 12 h nighttime sleep deprivation and 24 h sleep rebound. Two-way repeated-measures ANOVA with Fisher’s least significant difference (LSD) test detected a significant age × time interaction (F_(47, 5280)_ = 4.227; p < 0.0001) during sleep rebound. Asterisks above a line indicate time points at which the sleep percentage recovered differs significantly between 5d and 30d *wt* flies, respectively. (**J**) Sleep recovered at three different time points after sleep deprivation for 30d *wt* compared to 5d flies. n = 55-57 for both groups. Two-way ANOVA with Sidak’s multiple comparisons is shown. (**K**) Sleep profile for 30d *wt* flies treated with 5 mM Spd compared to untreated for 3 consecutive days. (**L**) Normalized cumulative sleep loss during 12 h nighttime sleep deprivation and 24 h sleep rebound. Two-way repeated-measures ANOVA with Fisher’s least significant difference (LSD) test detected a significant treatment × time interaction (F_(47, 4465)_ = 7.168; p < 0.0001) during sleep rebound. Asterisks above a line indicate time points at which the sleep percentage recovered differs significantly between 30d 5 mM Spd-treated and untreated flies, respectively. (**M**) Sleep recovered at three different time points after sleep deprivation for 30d 5 mM Spd-treated compared to untreated flies. n = 47-50 for both groups. Two-way ANOVA with Sidak’s multiple comparisons is shown. *p < 0.05; **p < 0.01; ***p < 0.001; ns, not significant. Error bars: mean ± SEM.

Consistently, we previously showed that Spd-supplied flies display suppressed PreScale during early aging (Gupta et al., 2016). Taken together, the effects of 1xBRP and Spd-supplementation on early aging-associated sleep pattern changes and PreScale-type plasticity (Figures 4, S4, 5 and S5A-S5E) (Gupta et al., 2016) suggest that PreScale is at least partially responsible for early aging-associated sleep pattern changes. We suspect that, under Spd supplementation, the normal early aging-associated increase of PreScale and consequently behavioral changes might be no longer required, allowing for preserved memory and a delay of age-associated synaptic changes coupled with an alleviation of daytime sleepiness. Thus, it is tempting to speculate that metabolic reprogramming by Spd supplementation via rejuvenating autophagy and mitochondrial respiration might suppress early aging-associated sleep patterns during early aging triggered by PreScale (Eisenberg et al., 2009; Gupta et al., 2013; Liang et al., 2021; Schroeder et al., 2021).

### Acute deep sleep reverses early aging-associated memory decline and PreScale-type plasticity

As mentioned earlier, in addition to sleep pattern changes (Figures 4, S4, 5 and S5), *Drosophila* aging is also associated with cognitive decline evident in reduced formation of aversive olfactory memories (Figures 6A and 6B) (Saitoe et al., 2005; Yamazaki et al., 2007; Gupta et al., 2013). Thus, we intended to further address the intersection of longevity, the ability of forming new memories, and early aging-associated sleep pattern changes, in conjunction with the role of PreScale plasticity. We argued that if PreScale-encoded additional sleep during early to mid-age might serve specific functions, and PreScale indeed would represent an acutely operating active mechanism to promote such changes, then triggering excessive deep sleep in mid-aged flies might allow to reset these age-associated behavioral and molecular regulations. To test this, we turned to the acute feeding of sleep-promoting Gaboxadol (4,5,6,7-Tetrahydroisoxazolo[5,4-c]pyridin-3-ol, THIP) which has been shown to rescue the formation of memories in standard memory mutants and Alzheimer’s disease model flies (Dissel et al., 2015; Dissel et al., 2017).

**Figure 6.**
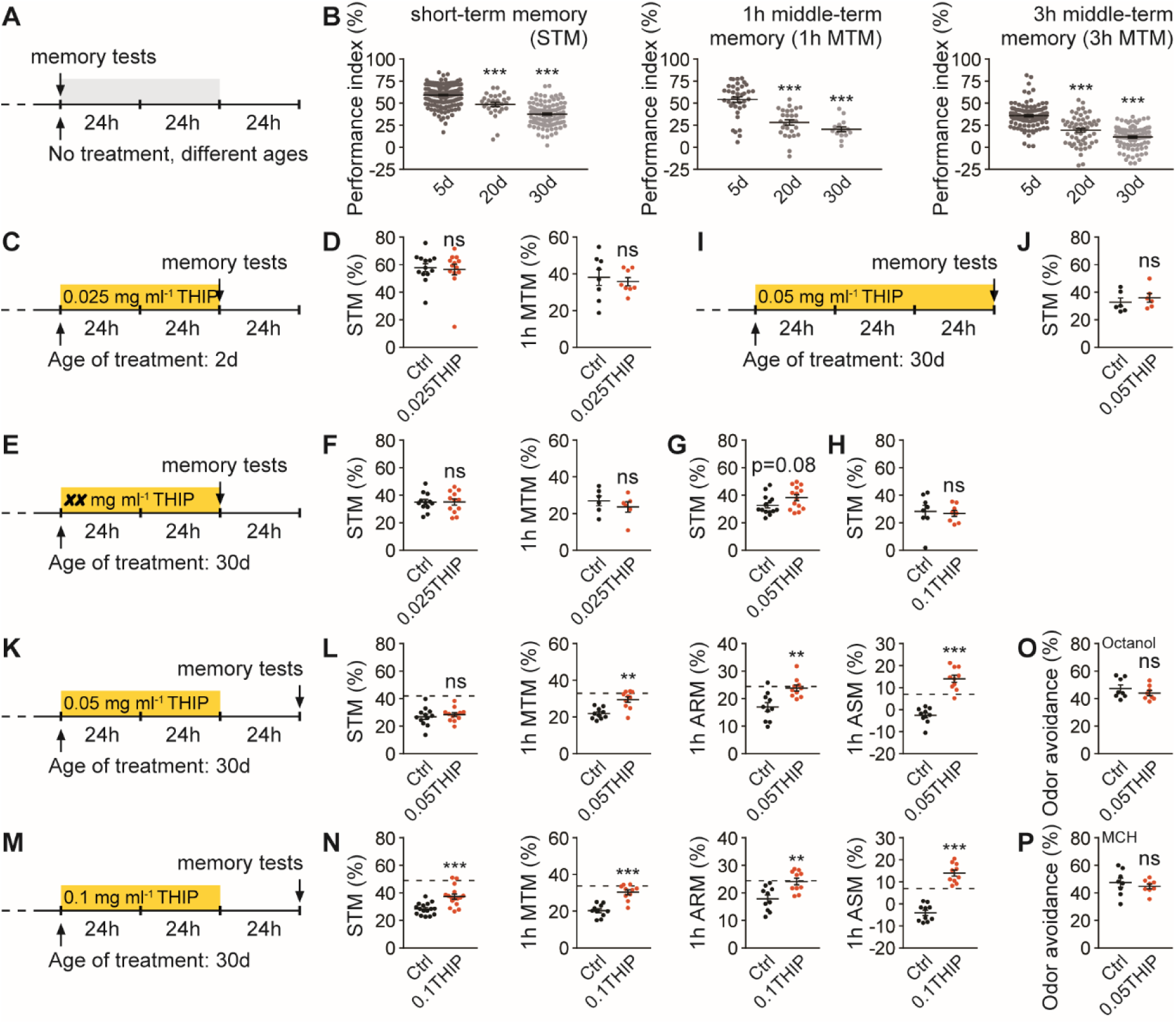
Acute deep sleep reverses age-associated memory decline. (**A** and **B**) Protocol for aging-related memory experiments (**A**) and typical age-associated memory decline in ages 20d and 30d *wt* flies (**B**), including short-term memory (STM), 1 hour middle-term memory (1h MTM) and 3h MTM. (**C** and **D**) Protocol (**C**) for 0.025 mg ml^-1^ THIP-treated *wt* flies at age 2d for 48 hours and tested thereafter for STM and 1h MTM immediately, compared to untreated siblings (**D**). n = 13 per group for STM, n = 8 per group for 1h MTM. (**E** and **F**) Protocol (**E**) for 0.025 mg ml^-1^ THIP-treated *wt* flies at age 30d for 48 hours and tested thereafter for STM and 1h MTM immediately, compared to untreated siblings (**F**). n = 12 per group for STM, n = 6 per group for 1h MTM. (**G** and **H**) With the protocol in Figure 6E, two higher THIP concentrations 0.05 mg ml^-1^ (**G**) and 0.1 mg ml^-1^ (**H**) were used to treat *wt* flies at age 30d for 48 hours and tested thereafter for STM immediately, compared to untreated siblings. n = 14 per group for 0.05 mg ml^-1^, n = 8 per group for 0.1 mg ml^-1^. (**I** and **J**) Protocol (**I**) for 0.05 mg ml^-1^ THIP-treated *wt* flies at age 30d for 72 hours and tested thereafter for STM immediately, compared to untreated siblings (**J**). n = 6 per group. (**K** and **L**) Protocol (**K**) for 0.05 mg ml^-1^ THIP-treated *wt* flies at age 30d for 48 hours and tested 24 hours after treatment for STM, 1h MTM, 1h anesthesia-resistant memory (ARM) and anesthesia-sensitive memory (ASM), compared to untreated siblings (**L**). n = 10-12 for all groups. (**M** and **N**) Protocol (**M**) for 0.1 mg ml^-1^ THIP-treated *wt* flies at age 30d for 48 hours and tested 24 hours after treatment for STM, 1h MTM, 1h ARM and 1h ASM, compared to untreated siblings (**N**). n = 16 per group for STM, n = 10 per group for 1h MTM, ARM and ASM. (**O** and **P**) With the protocol in Figure 6K except that 2d *wt* flies were used, odor avoidance was tested for the two odors (3-Octanol (**O**) and MCH (**P**), i.e., 4-methylcyclohexanol) used for olfactory memory experiments. n = 8 per group. Dashed lines indicate the average scores of respective memory components of 5d control *wt* flies without THIP treatment. Mann Whitney test is shown. *p < 0.05; **p < 0.01; ***p < 0.001; ns, not significant. Error bars: mean ± SEM.

Before testing memory with different protocols of THIP treatment, we first confirmed the sleep-promoting effects of THIP by feeding 5d young *wt* flies with both a low concentration of 0.025 mg ml^-1^ (0.025THIP) and a higher concentration of 0.1 mg ml^-1^ (0.1THIP) (Figures S6A-S6F) (Dissel et al., 2015). As feeding with 0.025THIP had already substantial sleep-promoting effects (Figures S6A-S6F), we first decided to feed young flies with 0.025THIP at age 2d for 2 days (48 hours) and measure olfactory memory immediately thereafter (Figure 6C). After training flies using paired associative aversive olfactory conditioning, the animals were tested either right away for short-term memory (STM) or 1 hour later for 1h middle-term memory (1h MTM). We did not find any differences in both STM and 1h MTM between 0.025THIP fed and unfed flies at age 2d (Figures 6C and 6D). It seems plausible that juvenile flies are still capable of forming memory at a maximum capacity and additional acute sleep might just not sufficient to boost olfactory memory formation. Thus, we continued by testing 30d flies, which displayed clearly reduced memories when compared to 5d young animals (Figures 6A and 6B).

However, feeding 30d *wt* flies with 0.025THIP for 48 hours followed by immediate testing for olfactory memory did not show any improvement for both STM and 1h MTM (Figures 6E and 6F). We wondered if the concentration 0.025THIP was insufficient to provoke any beneficial effects, and thus increased the concentrations to 0.05THIP and further to 0.1THIP. However, both concentrations did not provoke any significant improvement in STM (Figures 6E, 6G, 6H and S6G-S6K). It should be said, however, that when feeding 0.05THIP with this protocol, STM scores tended to increase (Figure 6G), though not significantly. Another obvious protocol was to increase the duration of THIP feeding to 3 days (72 hours) which, however, also did not improve STM (Figures 6I and 6J).

Given that THIP is extremely sleep-promoting, we decided to feed age 30d flies with 0.05THIP for 48 hours but tested memories only 24 hours after the end of THIP feeding (Figure 6K). Indeed, though STM was comparable to unfed control flies, 1h MTM was clearly improved over levels of unfed controls of the same age (Figures 6K and 6L). MTM is composed of two memory components which can be dissected through amnestic cooling, anesthesia-resistant memory (ARM) and anesthesia-sensitive memory (ASM) (Quinn and Dudai, 1976; Margulies et al., 2005). ARM is measured after amnestic cooling and ASM derives from subtracting ARM from MTM scores (Quinn and Dudai, 1976; Margulies et al., 2005). Notably, both ARM and ASM were fully restored by 0.05THIP treatment (Figures 6K and 6L). We further tested with a higher concentration of 0.1 mg ml^-1^ THIP (0.1THIP) (Figure 6M). This time, STM was also improved, and 1h MTM, ARM and ASM were all fully restored, just like under 0.05THIP treatment (Figures 6M and 6N). Importantly, THIP-feeding did not affect odor sensation, which is critical for olfactory memory formation (Figures 6O and 6P).

As shown above, a quasi linear BRP increase is associated with the occurrence of early aging-associated dFB activity reprogramming, sleep pattern changes, memory decline and longevity (Figures 1, 2, 3, 4 and 6). Furthermore, Spd supplementation suppresses both sleep pattern changes (Figures 5 and S5) and PreScale during early aging (Gupta et al., 2016), and promotes memory and longevity (Eisenberg et al., 2009; Gupta et al., 2016; Tain et al., 2020). We wondered if acute THIP-feeding, which rescued memory decline (Figures 6K-6N), might also suppress the early aging-associated PreScale indicated by elevated BRP levels (Figure 7A). Indeed, immuno-histological examination revealed a clear reduction of BRP levels in 0.1 mg ml^-1^ THIP-fed 30d flies when compared to unfed siblings (Figures 7B-7D). Notably, this was specific to age 30d flies, as BRP levels were not different in 0.1THIP-fed 5d brains from non-supplemented sibling controls (Figures 7B-7D).

**Figure 7.**
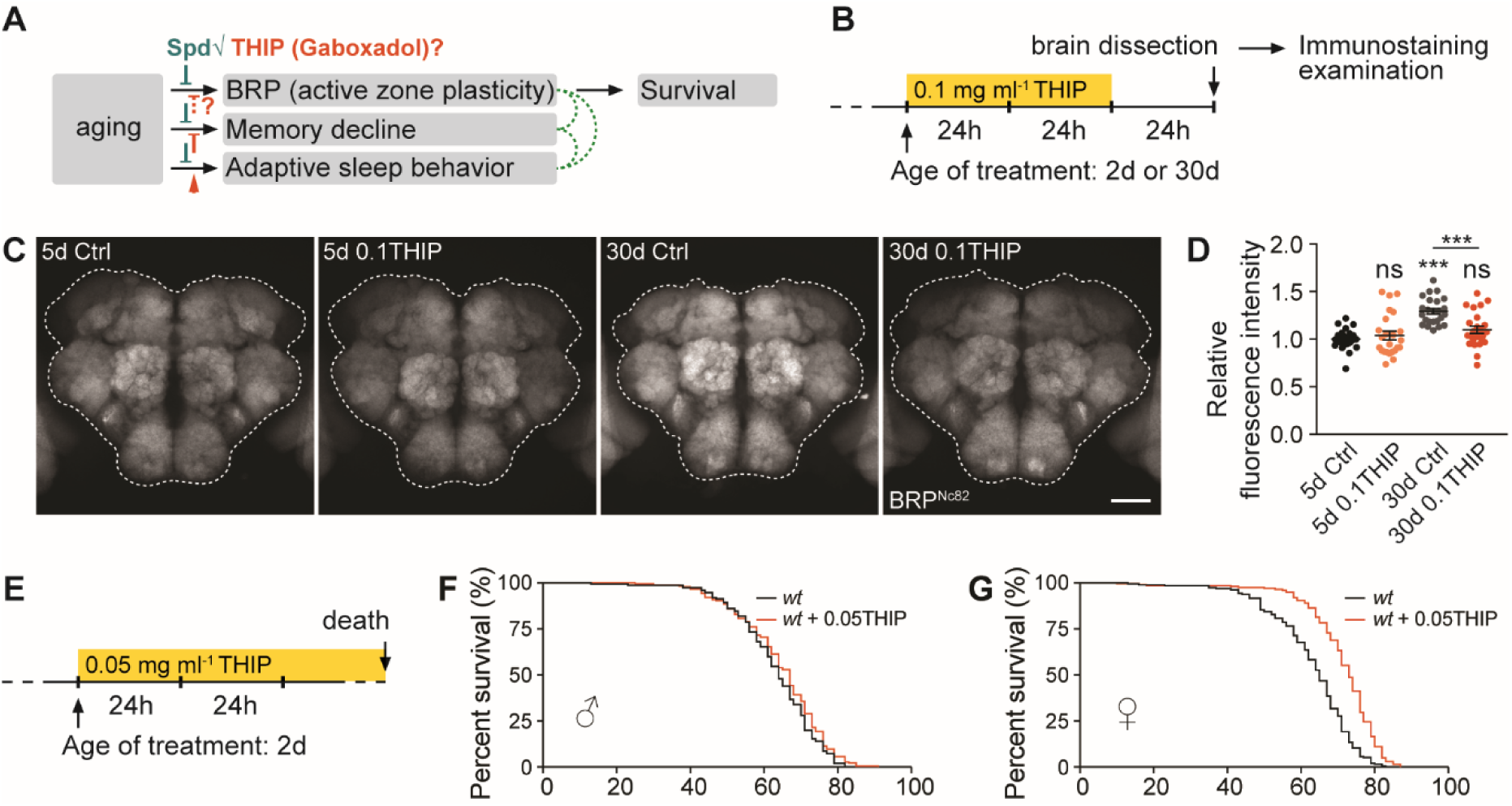
Excessive deep sleep reverses age-associated active zone plasticity/PreScale and boosts lifespan. (**A** and **B**) Rationale (**A**) and protocol (**B**) for immunochemistry examinations of fly brains of 0.1 mg ml^-1^ THIP-treated *wt* flies at ages 5d and 30d. (**C** and **D**) Confocal images (**C**) and whole-mount brain staining analysis (**D**) of BRP in *wt* female flies at ages 5d and 30d with or without 0.1 mg ml^-1^ THIP treatment. n = 24-25 for all groups. One-way ANOVA with Tukey’s post hoc tests is shown. Scale bar: 50 μm. (**E**) Protocol for longevity of 0.05 mg ml^-1^ THIP-treated *wt* flies. (**F** and **G**) Lifespan analysis of *wt* male and female flies treated with 0.05 mg ml^-1^ THIP. For male flies (**F**), n = 176 for treated compared to untreated (n = 150, ns). For female flies (**G**), n = 199 for treated compared to untreated (n = 192, p < 0.001). Gehan-Breslow-Wilcoxon test is shown. ***p < 0.001; ns, not significant. Error bars: mean ± SEM.

Motivated by the suppression of early aging-associated PreScale via THIP and Spd supplementation (Figures 7A-7D) (Gupta et al., 2016), and the longevity extension effect provoked by Spd supplementation (Eisenberg et al., 2009; Tain et al., 2020), we tested the lifespan of *wt* flies fed with 0.05 mg ml^-1^ THIP chronically at age 2d (Figure 7E). While *wt* male flies showed only a slight tendency towards lifespan extension (Figure 7F), female flies lived significantly longer upon 0.05THIP-feeding compared to unfed siblings (Figure 7G). Together with a suppression of age-associated memory decline and PreScale-type plasticity by acute THIP-feeding (Figures 6 and 7B-7D), it suggests that acute extensive deep sleep allowed for a reset/rejuvenation of a process underlying early brain aging, and promoted both lifespan and memory formation in mid-aged flies.

Taken together, we here used two distinct rejuvenation paradigms, THIP/Gaboxadol treatment and Spd supplementation, which provoked concordant effects: restoration of memory formation, increased longevity, and concomitantly an absence of PreScale in mid-aged flies (Figures 5, 6 and 7). Given the roles of PreScale in controlling early aging-associated sleep pattern changes and longevity (Figures 1, 3 and 4), this form of plasticity emerges at the intersection of sleep, longevity and age-associated memory decline. Notably, PreScale is also reversibly triggered by sleep deprivation for sleep homeostasis (Gilestro et al., 2009; Huang et al., 2020; Huang and Sigrist, 2020), suggesting a potential acute requirement of PreScale in resetting and denoising of circuits.

## DISCUSSION

Resilience designates the capacity of a system, enterprise or person to maintain its core purpose and integrity in the face of drastically changing circumstances (McEwen et al., 2015; McEwen, 2016). Resilience in human physiology is the capacity of adaptation in counteracting stress and adversity, while maintaining normal psychological and physical functions (Parrino and Vaudano, 2018). The brain as a central regulator of stress integration determines what is threatening, stores memories and regulates physiological adaptations across the aging trajectory (McEwen et al., 2015; McEwen, 2016; Aron et al., 2022). Thereby, sleep homeostasis is tightly linked to resilience mediated by the brain (McEwen et al., 2015; Parrino and Vaudano, 2018). Animal and patient data indicate that the impaired sleep observed to be associated with neurodegeneration is not only a result of an underlying circuit malfunction but might also directly contribute to neurodegeneration (Owen and Veasey, 2020). A key result of stress probably lies in the plastic reorganization of neural architecture and synaptic connections, which in turn may be a sign of successful adaptation (McEwen et al., 2015; Aron et al., 2022).

The aging process particularly challenges brain resilience (McEwen et al., 2015; McEwen, 2016; Aron et al., 2022). Typically, age-associated molecular and behavioral changes occur gradually with aging (Carey et al., 2006; Yamazaki et al., 2007; Gupta et al., 2013; Lopez-Otin et al., 2013). In particular, cognitive functions and sleep patterns change progressively with increasing age (Ohayon et al., 2004; Saitoe et al., 2005; Koh et al., 2006; Donlea et al., 2014; Metaxakis et al., 2014; Campos-Beltran and Marshall, 2021). For sleep, human studies reported diverse sleep alterations in old adults, including the difficulties to fall asleep, the inability to maintain sleep (insomnia) and daytime sleepiness (hypersomnia) (Pandi-Perumal et al., 2002). Similarly in flies, age-associated alterations of sleep patterns have been extensively reported, but might be sensitive to the exact chosen experimental conditions (Koh et al., 2006; Brown et al., 2014; Metaxakis et al., 2014; Vienne et al., 2016; Curran et al., 2019; Ly and Naidoo, 2019; Wiggin et al., 2020). However, the mechanisms mediating the age-associated sleep pattern changes are poorly understood. Specifically, we still miss critical information of how aging affects neuronal and synaptic organization and plasticity, how such changes intersect with age-associated alterations of sleep pattern and cognitive function, whether neuronal and synaptic changes might be causally involved in age-associated behavioral changes and resilience, and whether critical periods of resilience might exist across the aging trajectory. Brain resilience unnecessarily protects all these entities in a concordant fashion, and it is exactly the differentiation of causative from correlative or even adaptive and protective changes which is a fundamental but still unsolved problem.

Our previous studies described a shared molecular signature at central *Drosophila* brain synapses upon aging (Gupta et al., 2016) or sleep loss (Huang et al., 2020), which we collectively refer to as PreScale. In this study, we investigated the dynamic range and temporal profile of PreScale, and explored its consequences for cognitive fitness and sheer longevity. We provide evidence suggesting that this form of broadly distributed presynaptic plasticity provides a resilience module, which likely forms on-demand and allows the brain to cope with the consequences of aging and sleep deprivation. At the same time, PreScale might acutely prioritize tasks and establish trade-offs between survival and longevity on the one and memory formation on the other hand (Figure S7). These conclusions are based on the following new findings.

Firstly, PreScale builds up with age to peak at mid-age but at later advanced age declines again (Figure 1). Aging and sleep loss as two important stressors seemingly converge to trigger an at least very similar scenario of presynaptic active zone plasticity here. Notably, PreScale was reverted when mid-aged animals were pharmacologically triggered with extensive sleep for two days (Figures 7B-7D), and does not form in mid-aged animals when supplied with spermidine (Gupta et al., 2016).

Secondly, concerning resilience, when genetically mimicked to a moderate degree (3xBRP), PreScale promoted organismal longevity (Figures 3A-3E). It is certainly possible that chronic BRP titration triggered PreScale might in some regards differ from the presynaptic plasticity observed in either acutely sleep-deprived animals or in flies of mid-age. Still, direct comparisons indeed support that our experimental BRP titration can serve as a surrogate of early aging concerning behavioral alterations (Figures 2, 3A, 4, and S4). Importantly, 3xBRP efficiently promoted longevity in flies chronically challenged by sleep loss, but inefficient or even counterproductive in neurodegenerative conditions of other kind (Figures 3J-3M). Thus, 3xBRP might produce a kind of stress that regulates lifespan in a context-dependent manner (Figure 3B-3M). In contrast, genetically triggering very high, likely unphysiological BRP levels (4xBRP) robustly shortened lifespan (Figures 3A-3C and S3G-S3Q), arguing in direction of a “sweet spot” concerning its resilience-promoting function. Consistent with its dynamic range along the aging trajectory (Figure 3A), its critical period in relation to longevity promotion might indeed be the early aging period until mid-age. As we found BRP to promote sleep and regulate daily locomotion patterns in this early aging period (Figures 4 and S4), it is at least tempting to speculate that these presynaptic plasticity-driven processes might be causally related to resilience and extended longevity (Figure S7).

Thirdly, moderate BRP increases (3xBRP) tended to slightly reduce learning and memory capacity in the early aging period. Notably, high (4xBRP) levels even more severely reduce olfactory learning and memory (Gupta et al., 2016). The increased energy investment to support formation of longer-term memories operates in a trade-off relation between saving energy for better survival under food shortage or at advanced age (Placais and Preat, 2013; Placais et al., 2017). Similarly, early aging-associated sleep pattern changes, for example daytime sleepiness, could favor lower levels of energy consumption and consequently promote survival. PreScale might steer the trade-offs between the acute needs to entertain costly behaviors and/or to support sheer organismal fitness and longevity. Thus, we hypothesize that, during early aging, an optimal level of PreScale-type plasticity might be established at specific ages. It might be asked why our artificial increase of *brp* gene dose happens to be advantageous concerning longevity under our rearing conditions. Interestingly, learning ability can be substantially improved by artificial selection in animals ranging from *Drosophila* to rats (Burger et al., 2008), while *Drosophila* has been shown to make use of learned cues when tested in semi-natural environments (Zrelec et al., 2013). Along those lines, it will be very interesting to measure PreScale across the lifespan under different culture conditions, for example aging at lower temperatures (i.e., 18 °C compared to 25 °C) which were shown to extend lifespan and memory (Yamazaki et al., 2007).

PreScale might be a molecular/physiological manifestation of life strategy decisions balancing principal life-history related trade-offs. However, how would PreScale possible execute trade-offs between memory and longevity, given that PreScale itself is sleep-promoting? As healthy sleep supports normal learning and memory and also suggested by a previous study (Hendricks et al., 2001), it might be possible that genetic installation of PreScale promotes sleep which however does not support all functional aspects of natural sleep, leading to specific lifespan extension without benefiting cognitive functions. Considering the recently reported different sleep stages in flies (Yap et al., 2017; Tainton-Heap et al., 2021) and the sleep-supporting slow wave activities in the R5 network (Raccuglia et al., 2019), PreScale-encoded sleep need might define certain sleep stages which specifically promote lifespan but not support memory formation. Indeed, we recently showed that the short-term memory (STM) impairment of 4xBRP animals was fully rescued when BRP is eliminated specifically in the R5 neurons, while sleep was only partially suppressed (Huang et al., 2020). Thus, it is possible that such a manipulation allowed for a rebalance of the trade-offs which might reopen the window for executing the memory function of PreScale-encoded sleep, providing a plausible alternative explanation for early aging-associated memory impairment.

From our collective data, both Spd supplementation as well as triggering excessive sleep by THIP treatment allowed to attenuate PreScale in mid-aged flies (Figures 7A-7D and S7) (Gupta et al., 2016). We speculate that in these challenged brains some vital physiological cellular regulations might be no longer sufficiently plastic but instead steer into a detrimental direction. Notably, Spd supplementation age-protects mitochondrial respiration and autophagic flux, which normally decay in aging *Drosophila* brains (Liang et al., 2021; Schroeder et al., 2021). Efficient mitochondrial electron transport and autophagy in turn are coupled to sleep regulation (Kempf et al., 2019; Bedont et al., 2021). Moreover, mitochondrial functionality bidirectionally regulates early aging-associated PreScale-type plasticity revealed by BRP levels (Cho et al., 2021), and autophagy defects trigger PreScale in a non-cell autonomous manner (Bhukel et al., 2019). Thus, we speculate that Spd might delay the onset of early aging-associated sleep pattern changes and memory decline through its improvements of mitochondrial functions and autophagy and the consequent suppression of PreScale. Acute Gaboxadol/THIP treatment might equally reset the aging brain by reversing the early aging-associated decline in mitochondrial and autophagic functions (Bedont et al., 2021). It should be warranting to further dissect the molecular mechanisms that mediate the coupling between mitochondrial and autophagic status and the trade-offs of PreScale with early aging-associated sleep pattern changes and memory decline.

Concerning the mechanistic action of PreScale on synaptic and neurophysiological level, we observed a reprogramming of neuronal activity patterns of R5 neurons (Figures S2A and S2B) (Huang et al., 2020) and dFB neurons (Figures 2C and 2D), and the firing pattern and membrane excitability of the dFB neurons were changed upon genetic installation of PreScale and early aging (Figures 2E-2P). Indeed, a recent study suggests that dFB neurons promotes a specific form of sleep with brain activity that is different from spontaneous sleep (Tainton-Heap et al., 2021). As PreScale can *per se* change short-term plasticity and the transmission efficacy in a frequency dependent manner (Huang et al., 2020), we speculate that this form of synapse remodeling might promote the transmission of specific activity patterns. Favoring transmission in certain frequency domains might also explain its sleep-promoting action, as previous work identified slow wave oscillatory activities to be associated with sleep promotion via R5 neurons of the fly central complex (Raccuglia et al., 2019). In addition to R5 neurons, the interconnected dFB neurons have been shown to exhibit oscillatory activity and regulate both sleep and memory (Donlea et al., 2011; Dissel et al., 2015; Dissel et al., 2017; Yap et al., 2017; Tainton-Heap et al., 2021). It is tempting to speculate that dFB neurons might be at the intersection of PreScale-mediated trade-offs between longevity and cognitive functions, and the reduced neuronal activity and membrane excitability might explain the impaired memory in 3xBRP and 4xBRP animals. Potentially in the dFB/R5 network, skewing the representation of sensory information might also explain the reduced proclivity of flies under PreScale for forming new memories (Figure S7). How exactly the distinct effects of PreScale integrate from synapse over circuit and neuronal activity to behavioral level remains a warranting question for future analysis.

## STAR★METHODS

### Fly genetics and maintenance

*w^1118^* (*iso31*, BDSC#5905) was used as wildtype (*wt*), 2xBRP and background control. A *brp* null mutation (*c04298*, BDSC#85966) was utilized to reduce *brp* gene copy (Fouquet et al., 2009; Huang et al., 2020). A genomic *brp* P[acman] construct (Matkovic et al., 2013), which was mapped to be integrated into the 5’ UTR of *CG11357* (Figures S3G and S3H) (Potter and Luo, 2010), was used to increase *brp* gene copies from 2 to 3 and 4 (Huang et al., 2020), or to rescue the sleep phenotypes of 1xBRP flies (gBRP). A line with a P-element inserted at the 5’ UTR of *CG11357* (*EY12484*, BDSC#20838) was acquired to mimic the effects of a P-element at *CG11357* 5’ UTR, which was largely comparable to *wt* (Figures S3G-S3Q). Short sleep mutants *wake* (*EY02219*, BDSC#15858) and *inc* (*f00285*, BDSC#18307), and autophagy *atg7* mutant (*d06996*, BDSC#19257) were described previously (Bartlett et al., 2011; Stavropoulos and Young, 2011; Liu et al., 2014). Alzheimer’s disease model flies were generated by pan-neuronal *elav-Gal4*-driven expression of *Aβ-arctic* (BDSC#458 and #33774). *UAS-mCD8.GFP* (BDSC#32188) was expressed by *elav-Gal4* as control for Alzheimer’s disease model flies, or by *R23E10-Gal4* for patch-clamp electrophysiology. CaLexA (BDSC#66542) was expressed pan-neuronally by *elav-Gal4*, or in either dFB neurons marked by *R23E10-Gal4* (Pimentel et al., 2016) or R5 neurons driven by *R58H05-Gal4* (Liu et al., 2016). All fly strains were backcrossed to *w^1118^ wt* background for at least six generations to remove potential genetic modifiers.

Flies were raised under standard laboratory conditions on semi-defined medium (Bloomington recipe) under 12/12 h light/dark cycles with 65% humidity at 25 °C. Unless specifically stated, female flies at certain ages were used except aversive olfactory memory experiments in which mixed populations of both sexes were used.

## METHOD DETAILS

### Aging and longevity

During the development at larval stages, the density of larvae was purposely controlled across groups and genotypes. Aging was performed with mixed populations of male and female flies. To avoid stressing the flies during aging (for example flies step on each other), the amount of flies within each individual vials was grossly controlled across groups, genotypes and experiments. The aging flies were routinely flipped onto fresh food every second day or over the weekends until a desired age was reached for experiments.

Longevity experiments were carried out similarly, except that female and male flies were separated and sorted into a population of 20∼25 flies per vial/replicate at age 2d after fully mating. To reduce the variability in lifespan, a few different cohorts of flies were used for each experiment. Similar to aging flies, the longevity flies were regularly transferred onto fresh food and the amount of dead flies in each vial was recorded at each time of transfer until the death of the last fly of a replicate.

### Western blotting

Western blot analysis was carried out as previously reported (Huang et al., 2020). Basically, female flies at specific ages were dissected in ice-cold Ringer’s solution (pH = 7.3, 290-310 mOsm, with 5 mM HEPES-NaOH, 130 mM NaCl, 5 mM KCl, 2 mM MgCl_2_, 2 mM CaCl_2_ and 36 mM sucrose) between ZT6 and ZT10 (ZT, Zeitgeber Time). The brain samples were stored at -20 °C for short term (∼ 2 weeks) or at -80 °C for longer. Brain samples of the same experiment were fully homogenized by three rounds of intense vortexing in lysis buffer (0.5% Triton X-100, 2% SDS, 1x Protease inhibitor, 1x Sample buffer in PBS) followed by full-speed centrifugation for 6 min at 18 °C. The samples were then intensively vortexed to re-suspend the brain tissue and centrifuged again for 6 min at 18 °C. To avoid any saturation in the amount of proteins for detecting the differences, less than one brain’s supernatant was loaded into each lane for SDS-PAGE and immunoblotting. Thereafter, the blots were manually developed with ECL solutions and Kodak/GE films. The following primary antibodies were used: Mouse anti-BRP Nc82 (1:1000-1:2000), rabbit anti-BRP D2 (rb5228, 1:100000), guinea pig anti-Unc13A (14GP17, 1:2000), mouse anti-Dlg1 (4F3, 1:3000), mouse anti-Syn (3C11, 1:2000), mouse anti-Tubulin (Sigma T9026, 1:100000), mouse anti-Syx (8C3, 1:2000) and rabbit anti-Atg8a (Ab109364, 1:1000). The developed films without saturation were scanned by an EPSON V330 scanner in 16-bit grayscale tiff format (Huang et al., 2020).

### Immunostaining

Immunostaining was performed exactly as previously described (Huang et al., 2020). Adult female flies at certain ages or after THIP treatment were dissected in ice-cold Ringer’s solution and immediately fixed in 4% paraformaldehyde (PFA, pH = ∼7.3) for 30 min at room temperature. After fixing, brains were washed in 0.7% PBST (PBS with 0.7% Triton X-100, v/v) for 3 or 4 times for a total of 1 h and blocked in 0.7% PBST with 10% normal goat serum (NGS; v/v) for at least 2 h at room temperature. Primary antibodies were diluted in 0.7% PBST with 5% NGS for primary antibody incubation at 4 °C. Afterwards, brains were washed again in 0.7% PBST for at least 4 times and then incubated with secondary antibodies diluted in 0.7% PBST with 5% NGS in darkness. Finally, after secondary antibody incubation, brains were washed for at least 4 times and mounted in Vectashield and stored at 4 °C in darkness for confocal microscopy. Primary and secondary antibodies were incubated over a night, except for BRP Nc82 primary antibody, which requires at least two nights’ incubation for proper staining quality. The following primary antibodies were used: Mouse anti-BRP Nc82 (1:50) and chicken anti-GFP (ab13970, 1:1500). Goat anti-rabbit Cy5 and Alexa Fluor 647, goat anti-mouse Alexa 488 and goat anti-chicken Alexa 488 were diluted at 1:300 for secondary antibody incubation.

### Confocal microscope image acquisition, processing and analysis

Whole-mount brain samples were scanned with a Leica TCS SP8 confocal microscope under oil objectives. All the parameters of the microscope were set to avoid saturation and kept constant throughout each experiment for the purpose of comparing fluorescence intensity. All images (confocal and Western blot images) were processed and analyzed in ImageJ (Fiji) software (https://fiji.sc/). For stacks of brain images, an average intensity projection was chosen and the intensity of the single projected image was analyzed by manually drawing a region of interest (ROI) to determine the gray value within this ROI. The values for each replicate were normalized to *wt* or control and different replicates were pooled after normalization.

### Sleep and sleep deprivation

Sleep and sleep deprivation experiments were performed as previously published (Huang et al., 2020). Briefly, locomotor walking activity and sleep were recorded by *Drosophila* Activity Monitors (DAM2) from Trikinetics Inc. (Waltham, MA) in 12/12 h light/dark cycles at 25 °C with 65% humidity. Single flies at specific age were individually housed in Trikinetics glass tubes (5 mm inner diameter and 65 mm length) which had 5% sucrose and 2% agar in one side of the tube. The locomotor walking activity counts were uploaded every 1 min and the data from the first day was excluded due to the entrainment to new environments. A period of immobility without locomotor walking activity counts lasting for at least 5 min was defined as sleep (Shaw et al., 2000). Sleep and activity were analyzed using the Sleep and Circadian Analysis MATLAB Program (SCAMP) (Donelson et al., 2012).

Sleep deprivation was performed exactly as previously described (Huang et al., 2020). The DAM2 monitors were fixed onto a Vortexer Mounting Plate (VMP, Trikinetics) on an Analog Multi-Tube Vortexer controlled by a Trikinetics LC4 light controller and an acquisition software for sleep deprivation configuration. Pulses of vortex lasting for 1.2 s were applied randomly with inter-pulse intervals between 0 s and 40 s to fully sleep deprive flies during night from ZT12 to ZT24. As female flies show strong and consistent early aging-associated alterations in sleep patterns, females were used for all sleep experiments, except for Figures S3A-S3E, within which male flies were used.

### Aversive olfactory memory

Associative aversive olfactory memory experiments were performed as previously described (Gupta et al., 2016; Huang et al., 2020). Two aversive odors 3-octanol (Oct) (1:100 dilution) and 4-methylcyclohexanol (MCH) (1:100 dilution) were used as olfactory cues (odors were diluted in mineral oil) and 120 V AC current electrical shocks served as a behavioral reinforcer. Briefly, during training, about 100 flies were sequentially exposed to the first odor (conditioned stimulus, CS+, MCH or Oct) paired with electrical shocks (unconditioned stimulus, US) for 60 s followed by 60 s rest and then the second odor (CS-, Oct or MCH) without electrical shocks for 60 s. During testing, these flies were exposed simultaneously to both odors (CS+ and CS-) and had 60 s to choose between the two odors. A reciprocal experiment in which the other of the two odors was paired with electrical shocks was performed simultaneously. A memory index was calculated as the number of flies which chose the CS-odor minus the number of flies which chose the CS+ odor divided by the total number of flies. The final memory index was averaged from the two memory indices from the reciprocal experiments.

Short -term memory (STM) was tested immediately after training, while 1 hour middle-term memory (1h MTM) and 3h MTM were tested 1 hour or 3 hours after training. For 1 hour anesthesia-resistant memory (ARM), one of the two 1h MTM components, flies received cold shock on ice for 90 s, 30 min after training. 1h ARM was measured 30 min after cold shock and anesthesia-sensitive memory (ASM) was calculated by subtracting ARM from MTM.

Odor avoidance experiments were carried out similarly. Basically, naive flies were exposed to only one of the two odors and were allowed to choose between this odor and the air. The performance index was calculated as the number of flies which chose the air minus the number of flies which chose the odor divided by the total number of flies.

### Spermidine and THIP/Gaboxadol treatment

For acute THIP (Sigma-Aldrich, Cat# T101) treatment for memory tests, adult flies at certain ages were transferred onto normal fly food with specific concentrations of THIP at ∼ZT1 for either 48 hours or 72 hours. Flies were either tested immediately after treatment, or flipped back to normal fly food without THIP and tested 24 hours after THIP treatment. For acute THIP treatment for sleep experiments, specific concentrations of THIP were contained in 5% sucrose and 2% agar medium and sleep was measured as described above. For acute THIP treatment for immunostaining, flies were treated with 0.1 mg ml^-1^ THIP for 48 hours and dissected 24 hours after treatment.

Spermidine (Spd, Sigma-Aldrich, Cat# S2626) treatment has been described previously (Gupta et al., 2013; Gupta et al., 2016). Flies were raised and aged to specific days on 5 mM Spd containing normal fly food for sleep experiments with 5% sucrose and 2% agar medium.

### *In vivo* patch-clamp

5 and 20-day-old female flies (*R23E10>mCD8-GFP*) were used for *in vivo* patch-clamp electrophysiology recording of dFB neurons. Adult *Drosophila* preparation for *in vivo* electrophysiology recording was performed based on a former study (Murthy and Turner, 2013). Flies were anesthetized by keeping them on ice for 1-2 min and single flies were then fixed to the chamber with paraffin wax and dissected in external solution (103 mM NaCl, 3 mM KCl, 1.5 mM CaCl_2_, 4 mM MgCl_2_, 1 mM NaH_2_PO_4_, 26 mM NaHCO_3_, 5 mM TES (N-tris[hydroxymethyl]methyl-2-aminoethane sulfonic acid), 8 mM Trehalose, 10 mM Glucose, and 7 mM Sucrose, Osm = 280 ± 3) pre-equilibrated with carbogen (95% O_2_ / 5% CO_2_). The head cuticle on the posterior surface was peeled off to reveal the dFB neurons somata, and the glial sheath around the targeted area was focally removed with sharpened forceps. (Pimentel et al., 2016). GFP labeled dFB neurons were visualized under a 40 × objective. Signals were captured with a CCD digital camera (HAMAMATSU, ORCA-ER) mounted on the microscope.

Recording from dFB neurons was performed as described previously (Pimentel et al., 2016). During recording, the preparation was superfused continuously with the external solution. Somata of dFB neurons were targeted with pipettes (8-10 MΩ) filled with internal solution (140 mM K-aspartate, 1 mM KCl, 10 mM HEPES, 1 mM EGTA, 4 mM MgATP and 0.5 mM Na_3_GTP, pH = 7.3 and Osm = 265). Recordings were acquired with a MultiClamp 700B amplifier (Molecular Devices) and sampled with a Digidata 1440A interface (Molecular Devices). Signals were filtered at 6 kHz and digitized at 10-20 kHz. Data were analyzed using Clampfit 10.7 (Molecular Devices).

To measure the input resistance, small steps of 1 s hyperpolarizing current pulses were injected into the patched cell. To estimate the excitability of dFB neurons, cells were pre-hold at -60 mV followed by injection of a series of current steps (from -10 to 100 pA, 5 pA increment per step). Liquid junction potential of 13 mV was corrected off-line. Spikes were searched with the event detection function. The following criteria were used for bursts identification: inter-spike interval < 80 ms, number of spikes > 4. For comparisons of inter-spike interval distribution, Kolmogorov-Smirnov tests were performed. Cells with an access resistance higher than 50 MΩ were excluded from the analysis.

### Statistics

GraphPad Prism 7 was used to create the figures and perform most statistics. The Mann-Whitney test was used for comparison between two groups and one-way ANOVA with Tukey’s post hoc tests were used for multiple comparisons between multiple groups (≥3) with one factor. Two-way repeated-measures ANOVA was used for experiments with two factors. Longevity data was analyzed by Gehan-Breslow-Wilcoxon test in order to put more weight on earlier time points to compare differences between survival curves. Asterisks or ns (not significant) above a group indicate the comparison of this group to 2xBRP *wt* or controls; asterisks or ns above a line denote the comparison between the two specific groups for most statistical analyses, unless stated otherwise.

## DATA AND CODE AVAILABILITY

Original/source data are available from the corresponding author upon reasonable request.

## ACKNOWLEDGMENTS

We thank S. Yildirim-Brochno, D. Wachmann and Y. Ni for technical support; the Bloomington Stock Center (BDSC) for fly lines; and the Developmental Studies Hybridoma Bank (DSHB) for antibodies. This work was supported by grants from the Deutsche Forschungsgemeinschaft (DFG; German Research Foundation) to S.J.S. (SFB1315 TP A08 and FOR2705 TP05). S.H. was supported by the Chinese Scholarship Council as well as by the Leibniz SAW SyMetAge.

## AUTHOR CONTRIBUTIONS

Conceptualization, S.H. and S.J.S.; Investigation, S.H., C.P. and C.B.B.; Writing, S.H. and S.J.S.

---

**Figure S1.**
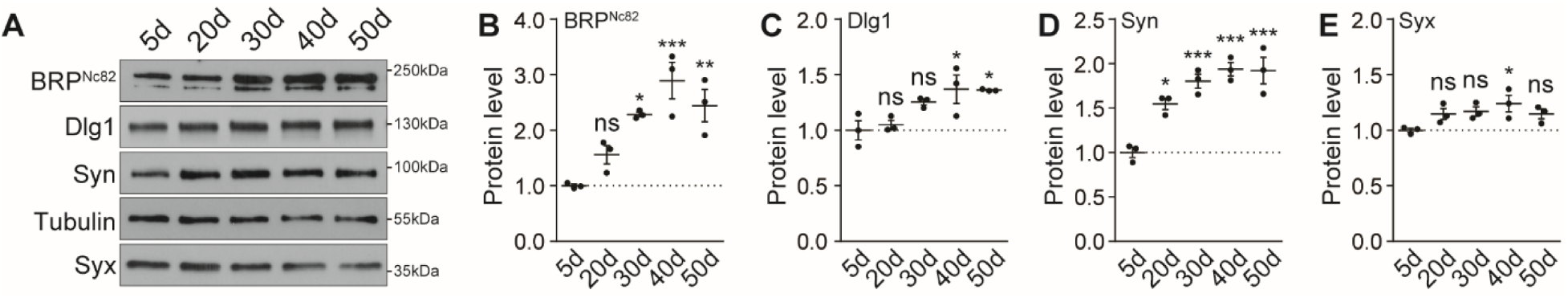
Synaptic plasticity with aging. (**A**-**E**) Representative Western blots (**A**) and statistics of a spectrum of synaptic proteins, including BRP Nc82 (**B**), Dlg1 (**C**), Syn (**D**), and Syx (**E**) in *w^1118^ wt* flies with aging. n = 3. One-way ANOVA with Dunnett’s post hoc tests is shown. *p < 0.05; **p < 0.01; ***p < 0.001; ns, not significant. Error bars: mean ± SEM.

**Figure S2.**
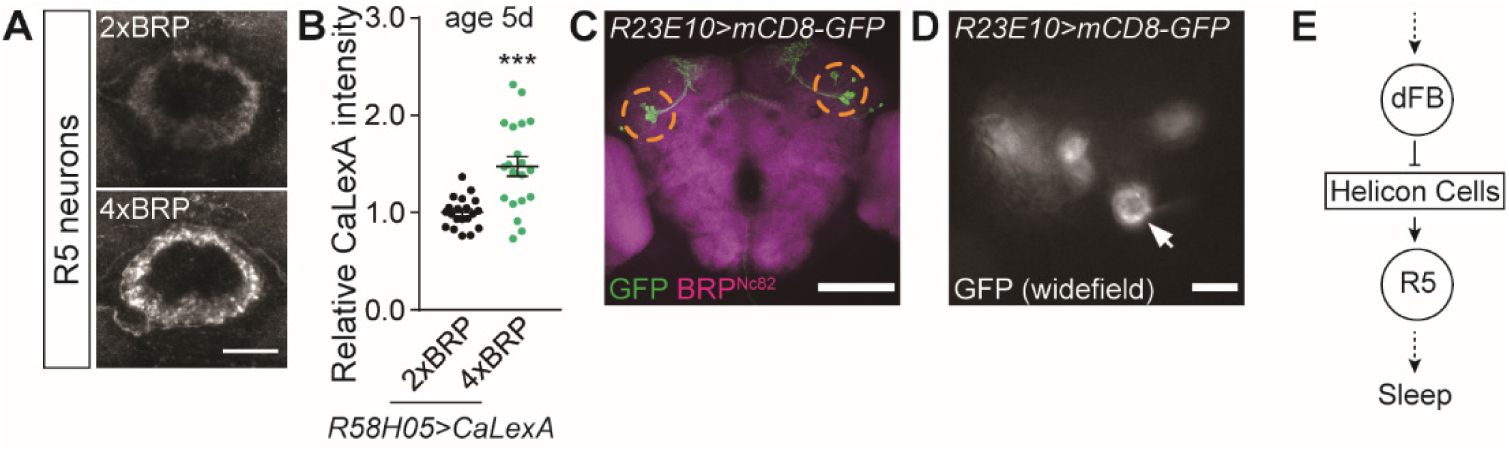
PreScale plasticity provokes activity reprogramming in the R5 neurons. (**A** and **B**) Confocal images (**A**) and whole-mount brain staining analysis (**B**) of CaLexA signal intensity with CaLexA expressed in R5 neurons by *R58H05-Gal4* in 2xBRP compared to 4xBRP flies. n = 20 for all groups. Scale bar: 20 μm. (**C**) Confocal image of the expression pattern of *R23E10-Gal4* indicated by GFP staining. Scale bar: 100 μm. (**D**) Widefield image of the cell bodies of *R23E10-Gal4* indicated by live GFP signal, the cell body localization is indicated in the dashed circles of (**C**). Scale bar: 10 μm. (**E**) Scheme of an interconnected sleep circuit in the central complex composed of the dorsal fan-shaped body (dFB, marked by *R23E10-Gal4*), Helicon cells and the ellipsoid body R5 neurons (R5 marked by *R58H05-Gal4*).

**Figure S3.**
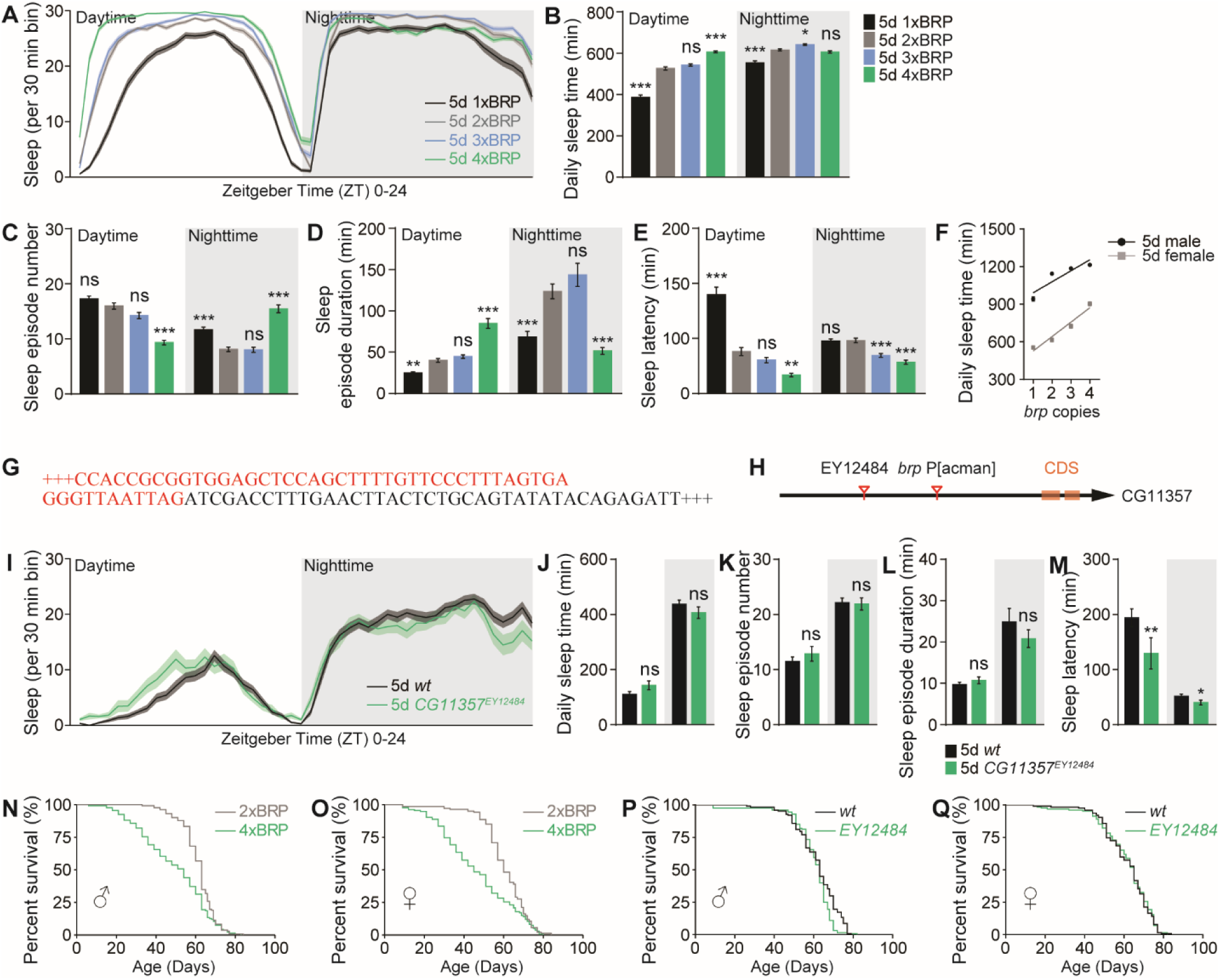
BRP promotes sleep in a dosage-dependent manner in male flies, similar to females. (**A**-**E**) Sleep structure of 1xBRP-4xBRP male flies averaged from measurements over 2-4 days, including sleep profile plotted in 30-min bins (**A**), daily sleep amount (**B**), number and duration of sleep episodes (**C** and **D**), and sleep latencies (**E**). n = 63–80. One-way ANOVA with Tukey’s post hoc tests is shown. (**F**) Linear regression analysis of daily sleep amount in flies with different *brp* copies in both male (R^2^ = 0.81) and female (R^2^ = 0.95). (**G**) Genomic mapping (Potter and Luo, 2010) and sequence of the integration site of the *brp* P[acman] transgenic construct. Red letters indicate *brp* P[acman] sequence and black letters indicate genomic *CG11357* gene sequence. (**H**) Simplified gene span of *CG11357* and the integration site of the *brp* P[acman] and another P-element mediated allele *EY12484* that are both localized at the 5’ UTR region of *CG11357*. CDS, coding DNA sequence. (**I**-**M**) Sleep structure of *EY12484* female flies averaged from measurements over 2-4 days, including sleep profile plotted in 30-min bins (**I**), daily sleep amount (**J**), number and duration of sleep episodes (**K** and **L**), and sleep latencies (**M**). n = 55 for *wt* control and n = 32 for *EY12484*. Mann-Whitney test is shown. (**N** and **O**) An independent experiment of the lifespan analysis of 2xBRP and 4xBRP flies. For male flies (**N**), n = 134 for 4xBRP compared to 2xBRP (n = 132, p < 0.001). For female flies (**O**), n = 134 for 4xBRP compared to 2xBRP *wt* control flies (n = 144, p < 0.001). (**P** and **Q**) Lifespan analysis of 2xBRP *wt* and *EY12484* flies. For male flies (**P**), n = 128 for *EY12484* compared to 2xBRP (n = 127, ns). For female flies (**Q**), n = 129 for *EY12484* compared to 2xBRP *wt* control flies (n = 127, ns). Gehan-Breslow-Wilcoxon test is shown for all longevity experiments. *p < 0.05; **p < 0.01; ***p < 0.001; ns, not significant. Error bars: mean ± SEM.

**Figure S4.**
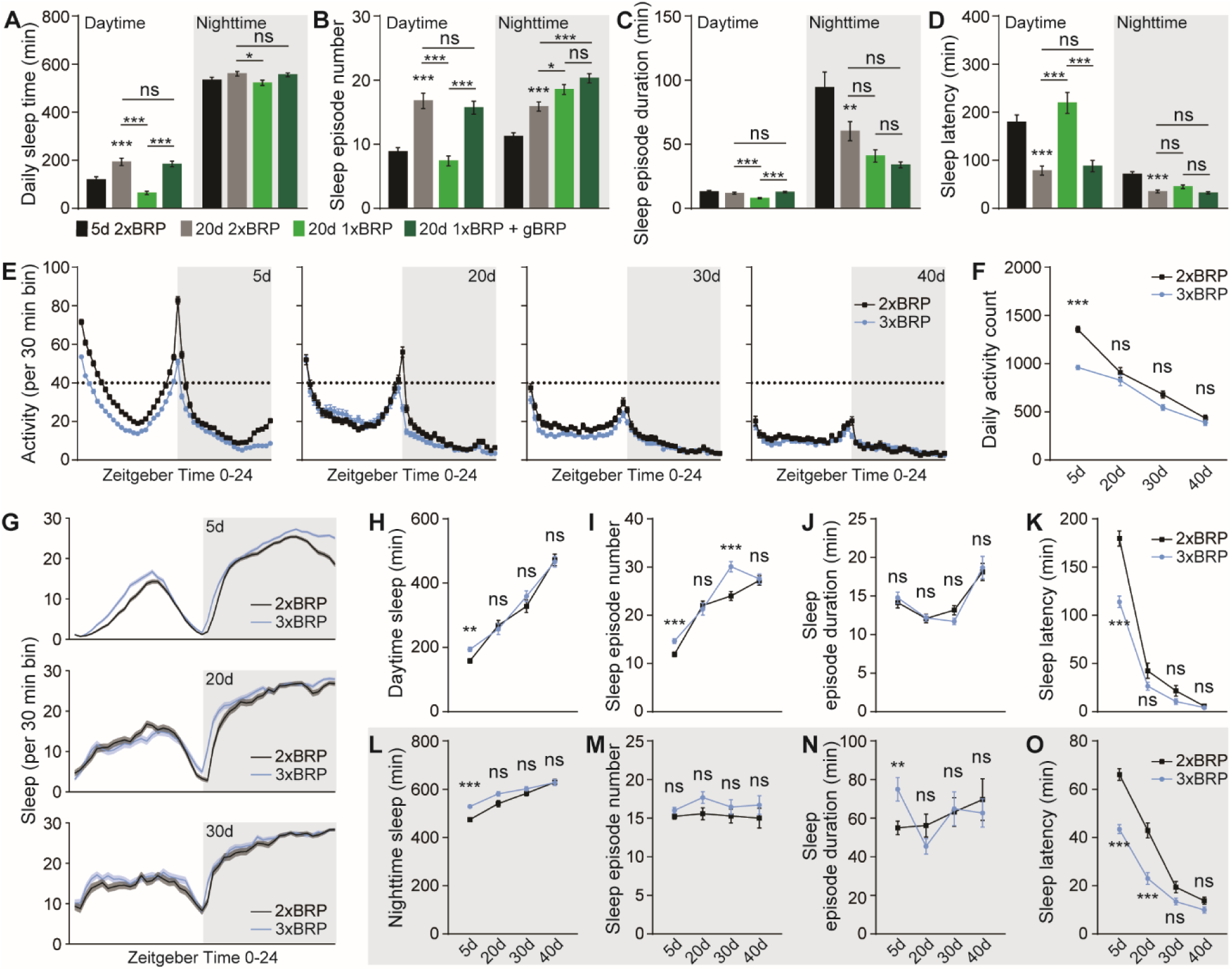
Early aging-associated sleep pattern changes of 3xBRP animals. (**A**-**D**) Sleep structure of 1xBRP flies at age 20d rescued by a transgenic *brp* copy (gBRP) and averaged from measurements over 2-3 days, including daily sleep amount (**A**), number and duration of sleep episodes (**B** and **C**), and sleep latencies (**D**). n = 63-64 for all groups. One-way ANOVA with Tukey’s post hoc tests is shown. (**E** and **F**) Locomotor walking activity distribution across the day (E) and averaged daily total walking activity (F) of 3xBRP compared to 2xBRP at ages 5d, 20d, 30d and 40d. (**G**-**O**) Sleep structure of 3xBRP flies at ages 5d, 20d, 30d and 40d averaged from measurements over 2-3 days, including sleep profile plotted in 30-min bins (**G**), daytime and nighttime sleep amount (**H** and **L**), number and duration of sleep episodes (**I**, **J**, **M** and **N**), and sleep latencies (**K** and **O**). n = 246-247 for 5d, n = 61-63 for 20d, n = 59-62 for 30d and n = 32 for 40d. Two-way ANOVA with Sidak’s multiple comparisons is shown. *p < 0.05; **p < 0.01; ***p < 0.001; ns, not significant. Error bars: mean ± SEM.

**Figure S5.**
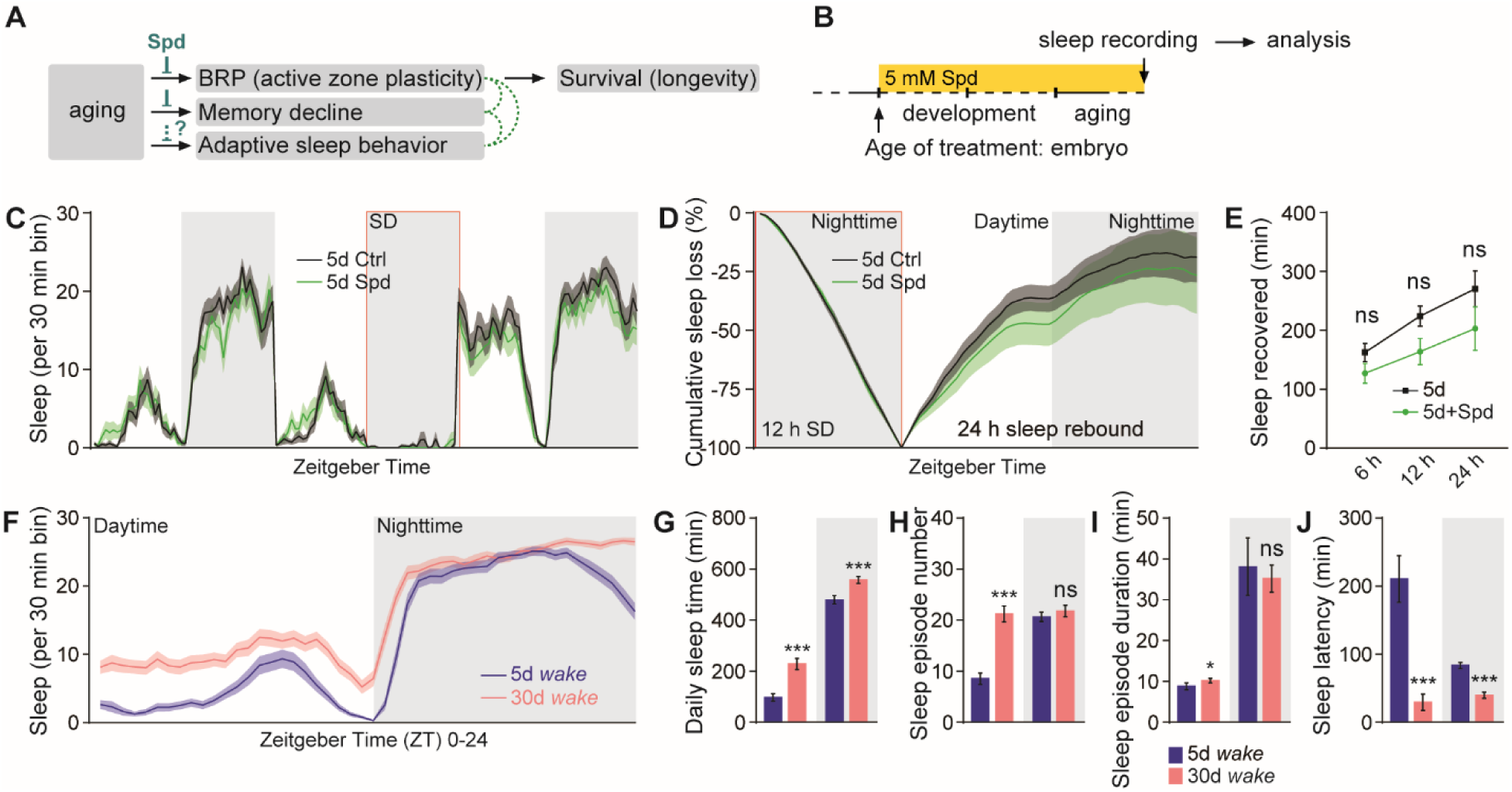
Early aging-associated alterations of sleep pattern in *wake* mutants. (**A** and **B**) Rationale (**A**) and protocol (**B**) for the consequence of Spd supplementation in age-associated alterations of sleep pattern. (**C**) Sleep profile for 5d *wt* flies treated with 5 mM Spd compared to untreated for 3 consecutive days. (**D**) Normalized cumulative sleep loss during 12 h nighttime sleep deprivation and 24 h sleep rebound. Two-way repeated-measures ANOVA with Fisher’s least significant difference (LSD) test did not detect any significant treatment × time interaction (F_(47, 2784)_ = 0.0038; p > 0.9999) during sleep rebound. (**E**) Sleep recovered at three different time points after sleep deprivation for 5d 5 mM Spd-treated compared to untreated flies. n = 29-31 for both groups. Two-way ANOVA with Sidak’s multiple comparisons is shown. (**F**-**J**) Sleep structure of 30d compared to 5d *wake* mutant flies from measurements over 2-3 days, including sleep profile plotted in 30-min bins (**F**), daily sleep amount (**G**), number and duration of sleep episodes (**H** and **I**), and sleep latencies (**J**). n = 47-48 for both groups. Mann Whitney test is shown. *p < 0.05; ***p < 0.001; ns, not significant. Error bars: mean ± SEM.

**Figure S6.**
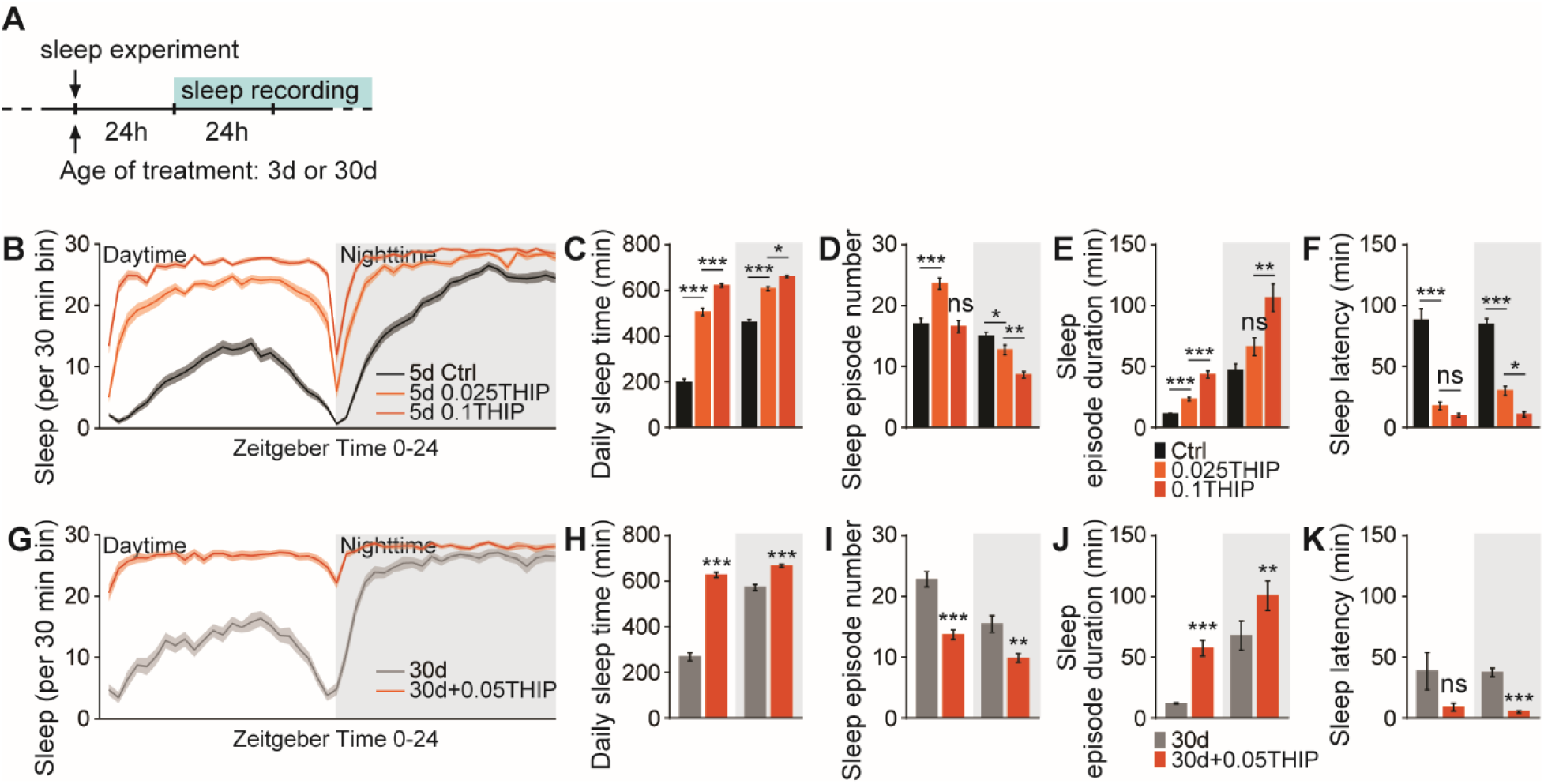
Acute deep sleep induced by THIP/Gaboxadol feeding. (**A**) Protocol for sleep test of *wt* flies treated with different concentrations of THIP at either age 3d or 30d. (**B**-**F**) Sleep structure of 5d *wt* flies fed with 0.05 mg ml^-1^ and 0.1 mg ml^-1^ THIP from measurements over 2-3 days, including sleep profile plotted in 30-min bins (**B**), daily sleep amount (**C**), number and duration of sleep episodes (**D** and **E**), and sleep latencies (**F**). n = 64 for untreated control *wt* flies, n = 32 for both 0.05 mg ml^-1^ and 0.1 mg ml^-1^ THIP-treated groups. One-way ANOVA with Tukey’s post hoc tests is shown. (**G**-**K**) Sleep structure of 30d *wt* flies fed with 0.05 mg ml^-1^ THIP from measurements over 2-3 days, including sleep profile plotted in 30-min bins (**G**), daily sleep amount (**H**), number and duration of sleep episodes (**I** and **J**), and sleep latencies (**K**). n = 31 for both groups. Mann Whitney test is shown. *p < 0.05; **p < 0.01; ***p < 0.001; ns, not significant. Error bars: mean ± SEM.

**Figure S7.**
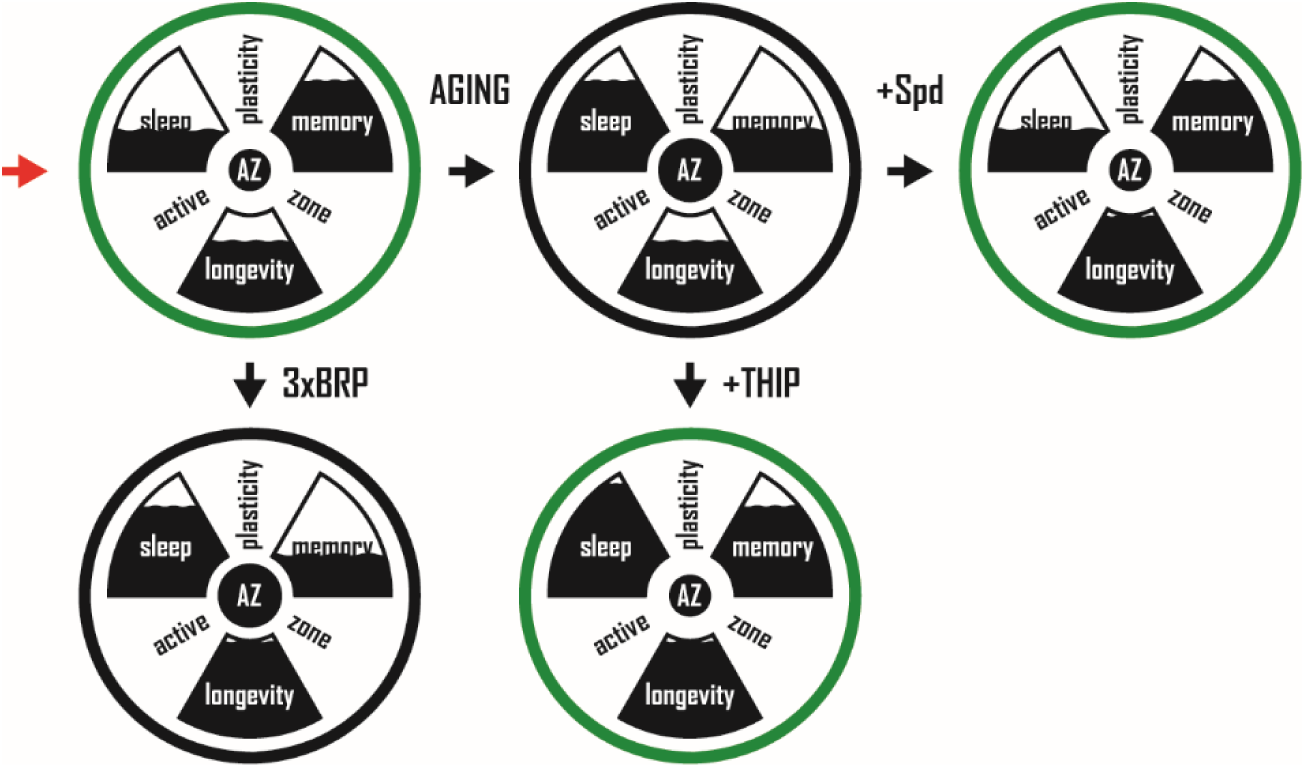
A model for presynaptic active zone plasticity (PreScale) in executing trade-offs between sleep, memory and longevity. A model for PreScale in executing trade-offs between sleep homeostasis, memory formation and longevity. Early aging provokes PreScale-type plasticity, which can be mimicked by titrating the gene copies of the master active zone (AZ) scaffold protein BRP. Two rejuvenation paradigms, spermidine (Spd) supplementation and THIP/Gaboxadol treatment, promote memory formation, extend longevity and suppress PreScale in mid-aged animals. Thus, PreScale likely executes behavioral adaptations and trade-offs during a still plastic phase of early brain aging.

## REFERENCES

Aron, L., Zullo, J., and Yankner, B.A. (2022). The adaptive aging brain. Current opinion in neurobiology 72, 91–100. Doi: 10.1016/j.conb.2021.09.009

Bartlett, B.J., Isakson, P., Lewerenz, J., Sanchez, H., Kotzebue, R.W., Cumming, R.C., Harris, G.L., Nezis, I.P., Schubert, D.R., Simonsen, A., et al. (2011). p62, Ref(2)P and ubiquitinated proteins are conserved markers of neuronal aging, aggregate formation and progressive autophagic defects. Autophagy 7, 572-583. Doi: 10.4161/auto.7.6.14943

Bedont, J.L., Toda, H., Shi, M., Park, C.H., Quake, C., Stein, C., Kolesnik, A., and Sehgal, A. (2021). Short and long sleeping mutants reveal links between sleep and macroautophagy. Elife 10. Doi: 10.7554/eLife.64140

Bhukel, A., Beuschel, C.B., Maglione, M., Lehmann, M., Juhasz, G., Madeo, F., and Sigrist, S.J. (2019). Autophagy within the mushroom body protects from synapse aging in a non-cell autonomous manner. Nature communications 10, 1318. Doi: 10.1038/s41467-019-09262-2

Bishop, N.A., Lu, T., and Yankner, B.A. (2010). Neural mechanisms of ageing and cognitive decline. Nature 464, 529–535. Doi: 10.1038/nature08983

Böhme, M.A., Beis, C., Reddy-Alla, S., Reynolds, E., Mampell, M.M., Grasskamp, A.T., Lutzkendorf, J., Bergeron, D.D., Driller, J.H., Babikir, H., et al. (2016). Active zone scaffolds differentially accumulate Unc13 isoforms to tune Ca(2+) channel-vesicle coupling. Nature neuroscience 19, 1311–1320. Doi: 10.1038/nn.4364

Brown, M.K., Chan, M.T., Zimmerman, J.E., Pack, A.I., Jackson, N.E., and Naidoo, N. (2014). Aging induced endoplasmic reticulum stress alters sleep and sleep homeostasis. Neurobiol Aging 35, 1431–1441. Doi: 10.1016/j.neurobiolaging.2013.12.005

Buckner, R.L. (2004). Memory and executive function in aging and AD: multiple factors that cause decline and reserve factors that compensate. Neuron 44, 195–208. Doi: 10.1016/j.neuron.2004.09.006

Burger, J.M., Kolss, M., Pont, J., and Kawecki, T.J. (2008). Learning ability and longevity: a symmetrical evolutionary trade-off in Drosophila. Evolution 62, 1294–1304. Doi: 10.1111/j.1558-5646.2008.00376.x

Campos-Beltran, D., and Marshall, L. (2021). Changes in sleep EEG with aging in humans and rodents. Pflugers Arch 473, 841–851. Doi: 10.1007/s00424-021-02545-y

Carey, J.R., Papadopoulos, N., Kouloussis, N., Katsoyannos, B., Muller, H.G., Wang, J.L., and Tseng, Y.K. (2006). Age-specific and lifetime behavior patterns in Drosophila melanogaster and the Mediterranean fruit fly, Ceratitis capitata. Exp Gerontol 41, 93–97. Doi: 10.1016/j.exger.2005.09.014

Cho, Y.H., Kim, G.H., and Park, J.J. (2021). Mitochondrial aconitase 1 regulates age-related memory impairment via autophagy/mitophagy-mediated neural plasticity in middle-aged flies. Aging Cell 20, e13520. Doi: 10.1111/acel.13520

Curran, J.A., Buhl, E., Tsaneva-Atanasova, K., and Hodge, J.J.L. (2019). Age-dependent changes in clock neuron structural plasticity and excitability are associated with a decrease in circadian output behavior and sleep. Neurobiol Aging 77, 158–168. Doi: 10.1016/j.neurobiolaging.2019.01.025

Davis, R.L. (2011). Traces of Drosophila memory. Neuron 70, 8–19. Doi: 10.1016/j.neuron.2011.03.012

Dissel, S. (2020). Drosophila as a Model to Study the Relationship Between Sleep, Plasticity, and Memory. Front Physiol 11, 533. Doi: 10.3389/fphys.2020.00533

Dissel, S., Angadi, V., Kirszenblat, L., Suzuki, Y., Donlea, J., Klose, M., Koch, Z., English, D., Winsky-Sommerer, R., van Swinderen, B., et al. (2015). Sleep restores behavioral plasticity to Drosophila mutants. Current biology : CB 25, 1270–1281. Doi: 10.1016/j.cub.2015.03.027

Dissel, S., Klose, M., Donlea, J., Cao, L., English, D., Winsky-Sommerer, R., van Swinderen, B., and Shaw, P.J. (2017). Enhanced sleep reverses memory deficits and underlying pathology in Drosophila models of Alzheimer’s disease. Neurobiol Sleep Circadian Rhythms 2, 15–26. Doi: 10.1016/j.nbscr.2016.09.001

Donelson, N.C., Kim, E.Z., Slawson, J.B., Vecsey, C.G., Huber, R., and Griffith, L.C. (2012). High-resolution positional tracking for long-term analysis of Drosophila sleep and locomotion using the "tracker" program. PLoS One 7, e37250. Doi: 10.1371/journal.pone.0037250

Donlea, J.M., Pimentel, D., Talbot, C.B., Kempf, A., Omoto, J.J., Hartenstein, V., and Miesenbock, G. (2018). Recurrent Circuitry for Balancing Sleep Need and Sleep. Neuron 97, 378–389 e374. Doi: 10.1016/j.neuron.2017.12.016

Donlea, J.M., Ramanan, N., Silverman, N., and Shaw, P.J. (2014). Genetic rescue of functional senescence in synaptic and behavioral plasticity. Sleep 37, 1427–1437. Doi: 10.5665/sleep.3988

Donlea, J.M., Thimgan, M.S., Suzuki, Y., Gottschalk, L., and Shaw, P.J. (2011). Inducing sleep by remote control facilitates memory consolidation in Drosophila. Science 332, 1571–1576. Doi: 10.1126/science.1202249

Eisenberg, T., Knauer, H., Schauer, A., Buttner, S., Ruckenstuhl, C., Carmona-Gutierrez, D., Ring, J., Schroeder, S., Magnes, C., Antonacci, L., et al. (2009). Induction of autophagy by spermidine promotes longevity. Nat Cell Biol 11, 1305–1314. Doi: 10.1038/ncb1975

Fouquet, W., Owald, D., Wichmann, C., Mertel, S., Depner, H., Dyba, M., Hallermann, S., Kittel, R.J., Eimer, S., and Sigrist, S.J. (2009). Maturation of active zone assembly by Drosophila Bruchpilot. The Journal of cell biology 186, 129–145. Doi: 10.1083/jcb.200812150

Gilestro, G.F., Tononi, G., and Cirelli, C. (2009). Widespread changes in synaptic markers as a function of sleep and wakefulness in Drosophila. Science 324, 109–112. Doi: 10.1126/science.1166673

Gill, S., Le, H.D., Melkani, G.C., and Panda, S. (2015). Time-restricted feeding attenuates age-related cardiac decline in Drosophila. Science 347, 1265–1269. Doi: 10.1126/science.1256682

Gupta, V.K., Pech, U., Bhukel, A., Fulterer, A., Ender, A., Mauermann, S.F., Andlauer, T.F., Antwi-Adjei, E., Beuschel, C., Thriene, K., et al. (2016). Spermidine Suppresses Age-Associated Memory Impairment by Preventing Adverse Increase of Presynaptic Active Zone Size and Release. PLoS Biol 14, e1002563. Doi: 10.1371/journal.pbio.1002563

Gupta, V.K., Scheunemann, L., Eisenberg, T., Mertel, S., Bhukel, A., Koemans, T.S., Kramer, J.M., Liu, K.S., Schroeder, S., Stunnenberg, H.G., et al. (2013). Restoring polyamines protects from age-induced memory impairment in an autophagy-dependent manner. Nature neuroscience 16, 1453–1460. Doi: 10.1038/nn.3512

Hendricks, J.C., Finn, S.M., Panckeri, K.A., Chavkin, J., Williams, J.A., Sehgal, A., and Pack, A.I. (2000). Rest in Drosophila is a sleep-like state. Neuron 25, 129–138. Doi:

Hendricks, J.C., Williams, J.A., Panckeri, K., Kirk, D., Tello, M., Yin, J.C., and Sehgal, A. (2001). A non-circadian role for cAMP signaling and CREB activity in Drosophila rest homeostasis. Nature neuroscience 4, 1108–1115. Doi: 10.1038/nn743

Huang, S., Piao, C., Beuschel, C.B., Gotz, T., and Sigrist, S.J. (2020). Presynaptic Active Zone Plasticity Encodes Sleep Need in Drosophila. Current biology : CB. Doi: 10.1016/j.cub.2020.01.019

Huang, S., and Sigrist, S.J. (2020). Presynaptic and postsynaptic long-term plasticity in sleep homeostasis. Current opinion in neurobiology 69, 1–10. Doi: 10.1016/j.conb.2020.11.010

Kaeberlein, M., Rabinovitch, P.S., and Martin, G.M. (2015). Healthy aging: The ultimate preventative medicine. Science 350, 1191–1193. Doi: 10.1126/science.aad3267

Kayser, M.S., Yue, Z., and Sehgal, A. (2014). A critical period of sleep for development of courtship circuitry and behavior in Drosophila. Science 344, 269–274. Doi: 10.1126/science.1250553

Keene, A.C., and Waddell, S. (2007). Drosophila olfactory memory: single genes to complex neural circuits. Nature reviews Neuroscience 8, 341–354. Doi: 10.1038/nrn2098

Kempf, A., Song, S.M., Talbot, C.B., and Miesenbock, G. (2019). A potassium channel beta-subunit couples mitochondrial electron transport to sleep. Nature 568, 230–234. Doi: 10.1038/s41586-019-1034-5

Kittel, R.J., Wichmann, C., Rasse, T.M., Fouquet, W., Schmidt, M., Schmid, A., Wagh, D.A., Pawlu, C., Kellner, R.R., Willig, K.I., et al. (2006). Bruchpilot promotes active zone assembly, Ca2+ channel clustering, and vesicle release. Science 312, 1051–1054. Doi: 10.1126/science.1126308

Klagges, B.R., Heimbeck, G., Godenschwege, T.A., Hofbauer, A., Pflugfelder, G.O., Reifegerste, R., Reisch, D., Schaupp, M., Buchner, S., and Buchner, E. (1996). Invertebrate synapsins: a single gene codes for several isoforms in Drosophila. The Journal of neuroscience : the official journal of the Society for Neuroscience 16, 3154–3165. Doi:

Koh, K., Evans, J.M., Hendricks, J.C., and Sehgal, A. (2006). A Drosophila model for age-associated changes in sleep:wake cycles. Proceedings of the National Academy of Sciences of the United States of America 103, 13843–13847. Doi: 10.1073/pnas.0605903103

Liang, Y., Piao, C., Beuschel, C.B., Toppe, D., Kollipara, L., Bogdanow, B., Maglione, M., Lutzkendorf, J., See, J.C.K., Huang, S., et al. (2021). eIF5A hypusination, boosted by dietary spermidine, protects from premature brain aging and mitochondrial dysfunction. Cell reports 35, 108941. Doi: 10.1016/j.celrep.2021.108941

Liu, S., Lamaze, A., Liu, Q., Tabuchi, M., Yang, Y., Fowler, M., Bharadwaj, R., Zhang, J., Bedont, J., Blackshaw, S., et al. (2014). WIDE AWAKE mediates the circadian timing of sleep onset. Neuron 82, 151–166. Doi: 10.1016/j.neuron.2014.01.040

Liu, S., Liu, Q., Tabuchi, M., and Wu, M.N. (2016). Sleep Drive Is Encoded by Neural Plastic Changes in a Dedicated Circuit. Cell 165, 1347–1360. Doi: 10.1016/j.cell.2016.04.013

Lopez-Otin, C., Blasco, M.A., Partridge, L., Serrano, M., and Kroemer, G. (2013). The hallmarks of aging. Cell 153, 1194–1217. Doi: 10.1016/j.cell.2013.05.039

Ly, S., and Naidoo, N. (2019). Loss of DmGluRA exacerbates age-related sleep disruption and reduces lifespan. Neurobiol Aging 80, 83–90. Doi: 10.1016/j.neurobiolaging.2019.04.004

Ly, S., Pack, A.I., and Naidoo, N. (2018). The neurobiological basis of sleep: Insights from Drosophila. Neurosci Biobehav Rev 87, 67–86. Doi: 10.1016/j.neubiorev.2018.01.015

Madeo, F., Eisenberg, T., Pietrocola, F., and Kroemer, G. (2018). Spermidine in health and disease. Science 359. Doi: 10.1126/science.aan2788

Mander, B.A., Rao, V., Lu, B., Saletin, J.M., Lindquist, J.R., Ancoli-Israel, S., Jagust, W., and Walker, M.P. (2013). Prefrontal atrophy, disrupted NREM slow waves and impaired hippocampal-dependent memory in aging. Nature neuroscience 16, 357–364. Doi: 10.1038/nn.3324

Mander, B.A., Winer, J.R., and Walker, M.P. (2017). Sleep and Human Aging. Neuron 94, 19–36. Doi: 10.1016/j.neuron.2017.02.004

Marck, A., Berthelot, G., Foulonneau, V., Marc, A., Antero-Jacquemin, J., Noirez, P., Bronikowski, A.M., Morgan, T.J., Garland, T., Jr., Carter, P.A., et al. (2017). Age-Related Changes in Locomotor Performance Reveal a Similar Pattern for Caenorhabditis elegans, Mus domesticus, Canis familiaris, Equus caballus, and Homo sapiens. J Gerontol A Biol Sci Med Sci 72, 455–463. Doi: 10.1093/gerona/glw136

Margulies, C., Tully, T., and Dubnau, J. (2005). Deconstructing memory in Drosophila. Current biology : CB 15, R700–713. Doi: 10.1016/j.cub.2005.08.024

Masuyama, K., Zhang, Y., Rao, Y., and Wang, J.W. (2012). Mapping neural circuits with activity-dependent nuclear import of a transcription factor. J Neurogenet 26, 89–102. Doi: 10.3109/01677063.2011.642910

Matkovic, T., Siebert, M., Knoche, E., Depner, H., Mertel, S., Owald, D., Schmidt, M., Thomas, U., Sickmann, A., Kamin, D., et al. (2013). The Bruchpilot cytomatrix determines the size of the readily releasable pool of synaptic vesicles. The Journal of cell biology 202, 667–683. Doi: 10.1083/jcb.201301072

McEwen, B.S. (2016). In pursuit of resilience: stress, epigenetics, and brain plasticity. Ann N Y Acad Sci 1373, 56–64. Doi: 10.1111/nyas.13020

McEwen, B.S., Bowles, N.P., Gray, J.D., Hill, M.N., Hunter, R.G., Karatsoreos, I.N., and Nasca, C. (2015). Mechanisms of stress in the brain. Nature neuroscience 18, 1353–1363. Doi: 10.1038/nn.4086

Metaxakis, A., Tain, L.S., Gronke, S., Hendrich, O., Hinze, Y., Birras, U., and Partridge, L. (2014). Lowered insulin signalling ameliorates age-related sleep fragmentation in Drosophila. PLoS Biol 12, e1001824. Doi: 10.1371/journal.pbio.1001824

Morrison, J.H., and Baxter, M.G. (2012). The ageing cortical synapse: hallmarks and implications for cognitive decline. Nature reviews Neuroscience 13, 240–250. Doi: 10.1038/nrn3200

Murthy, M., and Turner, G. (2013). Whole-cell in vivo patch-clamp recordings in the Drosophila brain. Cold Spring Harb Protoc 2013, 140–148. Doi: 10.1101/pdb.prot071704

Ohayon, M.M., Carskadon, M.A., Guilleminault, C., and Vitiello, M.V. (2004). Meta-analysis of quantitative sleep parameters from childhood to old age in healthy individuals: developing normative sleep values across the human lifespan. Sleep 27, 1255–1273. Doi: 10.1093/sleep/27.7.1255

Owen, J.E., and Veasey, S.C. (2020). Impact of sleep disturbances on neurodegeneration: Insight from studies in animal models. Neurobiol Dis 139, 104820. Doi: 10.1016/j.nbd.2020.104820

Pandi-Perumal, S.R., Seils, L.K., Kayumov, L., Ralph, M.R., Lowe, A., Moller, H., and Swaab, D.F. (2002). Senescence, sleep, and circadian rhythms. Ageing Res Rev 1, 559–604. Doi: 10.1016/s1568-1637(02)00014-4

Parrino, L., and Vaudano, A.E. (2018). The resilient brain and the guardians of sleep: New perspectives on old assumptions. Sleep Med Rev 39, 98–107. Doi: 10.1016/j.smrv.2017.08.003

Partridge, L., Deelen, J., and Slagboom, P.E. (2018). Facing up to the global challenges of ageing. Nature 561, 45–56. Doi: 10.1038/s41586-018-0457-8

Pfeiffenberger, C., and Allada, R. (2012). Cul3 and the BTB adaptor insomniac are key regulators of sleep homeostasis and a dopamine arousal pathway in Drosophila. PLoS Genet 8, e1003003. Doi: 10.1371/journal.pgen.1003003

Piao, C., and Sigrist, S.J. (2022). (M)Unc13s in Active Zone Diversity: A Drosophila Perspective. Frontiers in Synaptic Neuroscience 13. Doi: 10.3389/fnsyn.2021.798204

Pimentel, D., Donlea, J.M., Talbot, C.B., Song, S.M., Thurston, A.J., and Miesenbock, G. (2016). Operation of a homeostatic sleep switch. Nature 536, 333–337. Doi: 10.1038/nature19055

Placais, P.Y., de Tredern, E., Scheunemann, L., Trannoy, S., Goguel, V., Han, K.A., Isabel, G., and Preat, T. (2017). Upregulated energy metabolism in the Drosophila mushroom body is the trigger for long-term memory. Nature communications 8, 15510. Doi: 10.1038/ncomms15510

Placais, P.Y., and Preat, T. (2013). To favor survival under food shortage, the brain disables costly memory. Science 339, 440–442. Doi: 10.1126/science.1226018

Potter, C.J., and Luo, L. (2010). Splinkerette PCR for mapping transposable elements in Drosophila. PLoS One 5, e10168. Doi: 10.1371/journal.pone.0010168

Quinn, W.G., and Dudai, Y. (1976). Memory phases in Drosophila. Nature 262, 576–577. Doi: 10.1038/262576a0

Raccuglia, D., Huang, S., Ender, A., Heim, M.M., Laber, D., Suarez-Grimalt, R., Liotta, A., Sigrist, S.J., Geiger, J.R.P., and Owald, D. (2019). Network-Specific Synchronization of Electrical Slow-Wave Oscillations Regulates Sleep Drive in Drosophila. Current biology : CB. Doi: 10.1016/j.cub.2019.08.070

Saitoe, M., Horiuchi, J., Tamura, T., and Ito, N. (2005). Drosophila as a novel animal model for studying the genetics of age-related memory impairment. Rev Neurosci 16, 137–149. Doi:

Schroeder, S., Hofer, S.J., Zimmermann, A., Pechlaner, R., Dammbrueck, C., Pendl, T., Marcello, G.M., Pogatschnigg, V., Bergmann, M., Muller, M., et al. (2021). Dietary spermidine improves cognitive function. Cell reports 35, 108985. Doi: 10.1016/j.celrep.2021.108985

Schulze, K.L., Broadie, K., Perin, M.S., and Bellen, H.J. (1995). Genetic and electrophysiological studies of Drosophila syntaxin-1A demonstrate its role in nonneuronal secretion and neurotransmission. Cell 80, 311–320. Doi:

Seugnet, L., Suzuki, Y., Donlea, J.M., Gottschalk, L., and Shaw, P.J. (2011). Sleep deprivation during early-adult development results in long-lasting learning deficits in adult Drosophila. Sleep 34, 137–146. Doi:

Shaw, P.J., Cirelli, C., Greenspan, R.J., and Tononi, G. (2000). Correlates of sleep and waking in Drosophila melanogaster. Science 287, 1834–1837. Doi:

Simonsen, A., Cumming, R.C., Brech, A., Isakson, P., Schubert, D.R., and Finley, K.D. (2008). Promoting basal levels of autophagy in the nervous system enhances longevity and oxidant resistance in adult Drosophila. Autophagy 4, 176–184. Doi: 10.4161/auto.5269

Stavropoulos, N., and Young, M.W. (2011). insomniac and Cullin-3 regulate sleep and wakefulness in Drosophila. Neuron 72, 964–976. Doi: 10.1016/j.neuron.2011.12.003

Sudhof, T.C. (2012). The presynaptic active zone. Neuron 75, 11–25. Doi: 10.1016/j.neuron.2012.06.012

Tabuchi, M., Monaco, J.D., Duan, G., Bell, B., Liu, S., Liu, Q., Zhang, K., and Wu, M.N. (2018). Clock-Generated Temporal Codes Determine Synaptic Plasticity to Control Sleep. Cell 175, 1213–1227 e1218. Doi: 10.1016/j.cell.2018.09.016

Tain, L.S., Jain, C., Nespital, T., Froehlich, J., Hinze, Y., Gronke, S., and Partridge, L. (2020). Longevity in response to lowered insulin signaling requires glycine N-methyltransferase-dependent spermidine production. Aging Cell 19, e13043. Doi: 10.1111/acel.13043

Tainton-Heap, L.A.L., Kirszenblat, L.C., Notaras, E.T., Grabowska, M.J., Jeans, R., Feng, K., Shaw, P.J., and van Swinderen, B. (2021). A Paradoxical Kind of Sleep in Drosophila melanogaster. Current biology : CB 31, 578–590 e576. Doi: 10.1016/j.cub.2020.10.081

Ulgherait, M., Midoun, A.M., Park, S.J., Gatto, J.A., Tener, S.J., Siewert, J., Klickstein, N., Canman, J.C., Ja, W.W., and Shirasu-Hiza, M. (2021). Circadian autophagy drives iTRF-mediated longevity. Nature 598, 353–358. Doi: 10.1038/s41586-021-03934-0

Vienne, J., Spann, R., Guo, F., and Rosbash, M. (2016). Age-Related Reduction of Recovery Sleep and Arousal Threshold in Drosophila. Sleep 39, 1613–1624. Doi: 10.5665/sleep.6032

Wagh, D.A., Rasse, T.M., Asan, E., Hofbauer, A., Schwenkert, I., Durrbeck, H., Buchner, S., Dabauvalle, M.C., Schmidt, M., Qin, G., et al. (2006). Bruchpilot, a protein with homology to ELKS/CAST, is required for structural integrity and function of synaptic active zones in Drosophila. Neuron 49, 833–844. Doi: 10.1016/j.neuron.2006.02.008

Wang, M., Gamo, N.J., Yang, Y., Jin, L.E., Wang, X.J., Laubach, M., Mazer, J.A., Lee, D., and Arnsten, A.F. (2011). Neuronal basis of age-related working memory decline. Nature 476, 210–213. Doi: 10.1038/nature10243

Weiss, J.T., and Donlea, J.M. (2021). Sleep deprivation results in diverse patterns of synaptic scaling across the Drosophila mushroom bodies. Current biology : CB 31, 3248–3261 e3243. Doi: 10.1016/j.cub.2021.05.018

Wiggin, T.D., Goodwin, P.R., Donelson, N.C., Liu, C., Trinh, K., Sanyal, S., and Griffith, L.C. (2020). Covert sleep-related biological processes are revealed by probabilistic analysis in Drosophila. Proceedings of the National Academy of Sciences of the United States of America 117, 10024–10034. Doi: 10.1073/pnas.1917573117

Woods, D.F., and Bryant, P.J. (1991). The discs-large tumor suppressor gene of Drosophila encodes a guanylate kinase homolog localized at septate junctions. Cell 66, 451–464. Doi:

Yamazaki, D., Horiuchi, J., Nakagami, Y., Nagano, S., Tamura, T., and Saitoe, M. (2007). The Drosophila DCO mutation suppresses age-related memory impairment without affecting lifespan. Nature neuroscience 10, 478–484. Doi: 10.1038/nn1863

Yankner, B.A., Lu, T., and Loerch, P. (2008). The aging brain. Annu Rev Pathol 3, 41–66. Doi: 10.1146/annurev.pathmechdis.2.010506.092044

Yap, M.H.W., Grabowska, M.J., Rohrscheib, C., Jeans, R., Troup, M., Paulk, A.C., van Alphen, B., Shaw, P.J., and van Swinderen, B. (2017). Oscillatory brain activity in spontaneous and induced sleep stages in flies. Nature communications 8, 1815. Doi: 10.1038/s41467-017-02024-y

Zrelec, V., Zini, M., Guarino, S., Mermoud, J., Oppliger, J., Valtat, A., Zeender, V., and Kawecki, T.J. (2013). Drosophila rely on learning while foraging under semi-natural conditions. Ecol Evol 3, 4139–4148. Doi: 10.1002/ece3.783

